# Hybrid Adipocyte-Derived Exosome Nano Platform for Potent Chemo-Phototherapy in Targeted Hepatocellular Carcinoma

**DOI:** 10.1101/2023.11.14.567027

**Authors:** Xinying Liu, Jiaxin Zhang, Shunzhe Zheng, Meng Li, Wenqian Xu, Jianbin Shi, Ken-ichiro Kamei, Chutong Tian

**Author notes:** Corresponding authors. E-mail address (CT), (KK).

## Abstract

The high prevalence and severity of hepatocellular carcinoma (HCC) present a significant menace to human health. Despite the significant advancements in nanotechnology-driven antineoplastic agents, there remains a conspicuous gap in the development of targeted chemotherapeutic agents specifically designed for HCC. Consequently, there is an urgent need to explore potent drug delivery systems for effective HCC treatment. Here we have exploited the interplay between HCC and adipocyte to engineer a hybrid adipocyte-derived exosome platform, serving as a versatile vehicle to specifically target HCC and exsert potent antitumor effect. A lipid-like prodrug of docetaxel (DSTG) with a reactive oxygen species (ROS)-cleavable linker, and a lipid-conjugated photosensitizer (PPLA), spontaneously co-assemble into nanoparticles, functioning as the lipid cores of the hybrid exosomes (HEMPs and NEMPs). These nanoparticles are further encapsuled within adipocyte-derived exosome membranes, enhancing their affinity towards HCC cancer cells. As such, cancer cell uptakes of hybrid exosomes are increased up to 5.73-fold compared to lipid core nanoparticles. Our *in vitro* and *in vivo* experiments have demonstrated that HEMPs not only enhance the bioactivity of the prodrug and extend its circulation in the bloodstream but also effectively inhibit tumor growth by selectively targeting hepatocellular carcinoma tumor cells. Self-facilitated synergistic drug release subsequently promoting antitumor efficacy, inducing significant inhibition of tumor growth with minimal side effects. Our findings herald a promising direction for the development of targeted HCC therapeutics.

**Graphical abstract:** An adipocyte-derived exosome nanoplatform has been developed for the combined therapy of both chemotherapy and photodynamic therapy. The hybrid adipocyte-derived exosome nanoplatform integrates the advantage of targeted co-delivery and combination therapy, which can stimulate the release of drugs under the tumor microenvironment, effectively enhanced targeted co-delivery in hepatocellular carcinoma cells, prolonged blood circulation time and improve the therapeutic effect of hepatocellular carcinoma tumors model.

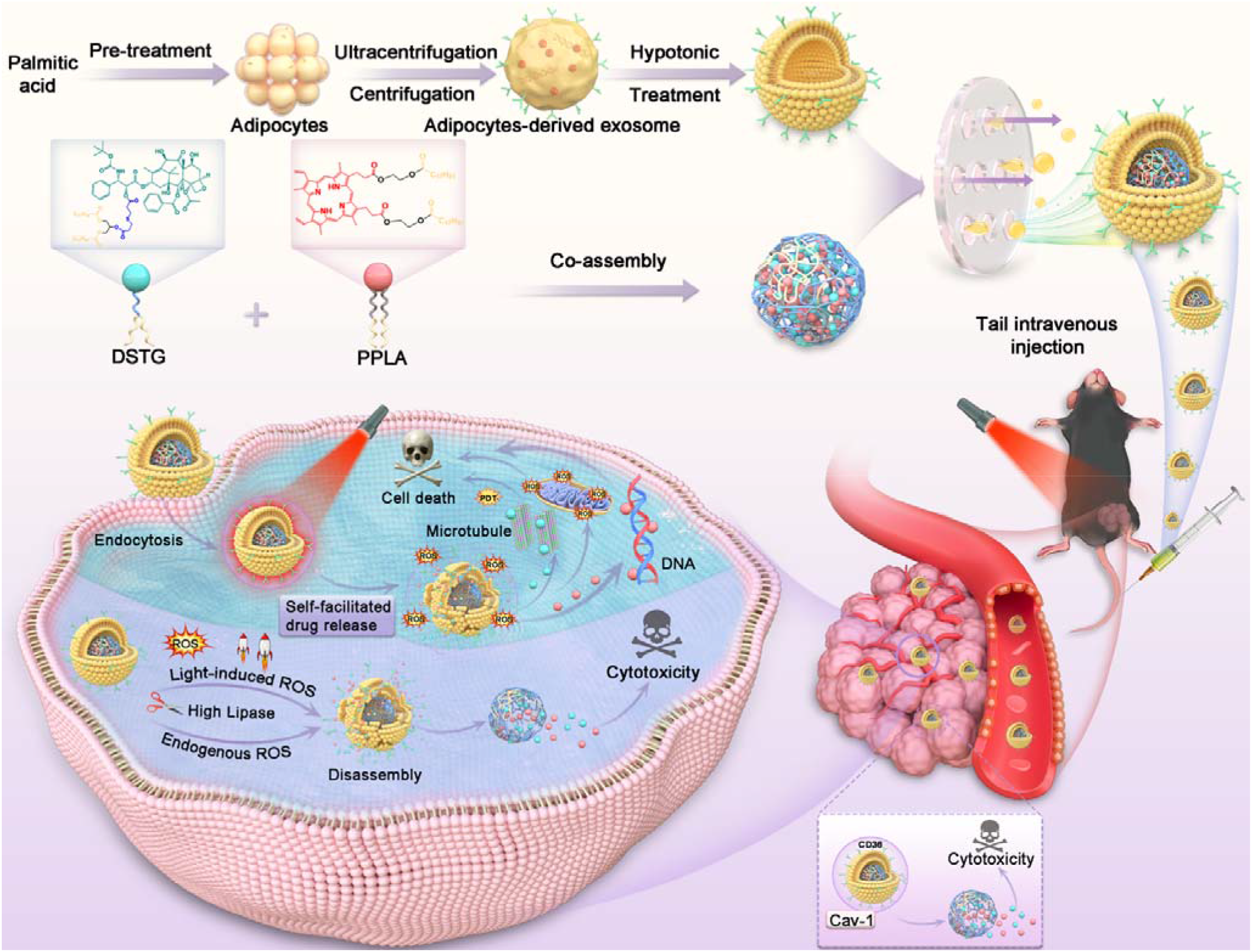

## Introduction

Hepatocellular carcinoma (HCC) is a prevalent primary cancer globally, often presenting with severe symptoms and leading to high recurrence rates and mortality. Historically, surgical resection of HCC and liver transplantation from compatible healthy donors have been the primary therapeutic interventions. However, these methods are hampered by the intricate surgical techniques required for resection and the challenge of finding compatible donors. Although alternative treatments like photodynamic therapy (PDT), chemotherapy, ablation therapy, and radiotherapy have been developed, effectively treating advanced HCC remains a formidable challenge due to systemic toxicity and severe adverse reactions.^1^ Traditional administration of solution-based of free therapeutic agents typically demonstrate limited selectivity in tumor targeting, suboptimal therapeutic synergy, and rapid clearance.^2^ Consequently, there is a pressing need to develop a novel, minimally invasive treatment for HCC patients.

Addressing these challenges, multi-modal therapy can exert synergistic anti-tumor effects, surpassing the additive and synergetic effects of standalone treatments.^3^ For instance, merging PDT with chemotherapy presents a potent anticancer approach. This strategy involves administering photosensitizers and chemotherapeutic agents, followed by targeted light activation to enhance therapeutic results.^4^ Despite the promise of multi-modal therapy, it continues to confront challenges, including variability of the pharmacokinetics and biodistribution of constituent drugs, concerns regarding circulatory, and the risk of inadvertent impacts on healthy tissues.

For advancement of multi-modal therapy, the employment of self-assembled prodrug nano-systems presents significant opportunities by offering improved pharmacokinetics relative to free drug forms, along with spatiotemporal control over drug activity.^5^ This strategy also enables high drug loading without the risks of excipient-related toxicity and allowing for targeted drug release within tumor via stimuli-responsive linkages.^6^ Furthermore, the choice of stimuli-responsive linkage ensures optimal drug release from the prodrug. Notably, the single thioether linkage exhibits pronounced responsiveness to reactive oxygen species (ROS), leading to swift drug release, spurred by the elevated ROS levels in tumor cells, but not healthy cells.^7^ Intriguingly, when subjected to light irradiation, the photosensitizer not only generates ROS, which can damage organelles, but also swiftly cleaves the ROS-activated linkage. This process amplifies the cytotoxic drug’s activity in responsive prodrugs, creating a “cascade release” mechanism conducive for effective cancer therapy.^8^ Nonetheless, this strategy still grapples with challenges in actively targeting tumor sites and uncertain biocompatibility, due to the fleeting interactions inherent in synthetic nano-prodrugs.

To address this challenge, we meticulously realized exosomes—nanometer-scale vesicles composed of a phospholipid bilayer^9^ secreted by cells. In fact, endogenous exosomes have been widely employed in nanodrug delivery systems owing to their unique advantages compared to exogenous nanomaterials. These advantages include optimal particle size, excellent biocompatibility, biodegradability, evasion of macrophage phagocytosis due to membrane-bound proteins, extended *in vivo* circulation, and precise targeting properties.^10^ Moreover, recent studies have indicated a heightened risk of developing HCC closely linked to obesity.^11^ In HCC, adipocyte-derived exosomes (ad-exos) play distinctive roles, facilitating the transport of lipids, proteins associated with fatty acid oxidation, and mRNA.^12^ This transport supports tumor proliferation, invasion, and metastasis. The integral role of ad-exos in this interplay offers a new frontier for targeted drug delivery against HCC. Drawing inspiration from these unique attributes, we hypothesize that ad-exos could be potent vehicles for drug delivery aimed at HCC, given their excellent biocompatibility and potential HCC targeting property.

Here we develop a novel hybrid adipocyte-derived exosome nano platform for potent chemo-phototherapy in targeting HCC, enabling effective tumor targeting, dual-responsive and self-amplifying cascade drug release mechanisms, and enhanced stability *in vivo*. To integrate self-assembled nano-prodrugs into lipid-rich adipocytes-derived exosomes (ad-exos), the hydrophobic side chains, such as polyunsaturated fatty acids (PUFAs) and vitamin E, are introduced to promote the spontaneous self-assembly of prodrugs into nanoparticles (NPs).^13^ In light of the lipid-rich features of ad-exos, hydrophobic triglycerides (TGs) are ideal as assembly blocks and also serve as gates responsive to lipase, ensuring a more site-specific drug release at tumor sites.^14^ Hence, we synthesize two lipidic prodrugs to encapsulate both chemotherapeutic and PDT agents into the nano-assembly, concurrently replacing the cores of ad-exos with this configuration: a thioether bond-bridged triglyceride-like prodrug of DTX (referred to DSTG) and a PUFAylation prodrug of protoporphyrin IX (PpIX) (termed PPLA). Remarkably, these two prodrugs are able to spontaneously co-assemble into nanoparticles in deionized water, eliminating the need of surfactants, and subsequently function as the lipid core of ad-exos. The resulting hybrid ad-exo camouflaged prodrug nanoplatforms are endowed with enhanced targeting efficiency towards HCC, self-amplified synergistic drug release capacity, retained biocompatibility and biometabolability under physiological conditions, consequently demonstrating significant anti-tumor activity. The innovate approach is poised to improve the efficacy of multi-modal therapy and broadening the scope of targeted HCC treatment.

## Results and discussion

### Adipocyte-Derived Exosomes Promote Liver Cell Uptake

To obtain the functional adipocyte-derived exosomes, 3T3-L1 mouse preadipocytes were induced into mature adipocytes using the “cocktail” differentiation method, with palmitic acid mimicking obesity conditions (Figure 1A). Successful adipocyte differentiation was confirmed through oil red O staining, revealing red-stained lipid droplets characteristic of mature adipocytes (Figure S1). Subsequently, monensin sodium (MON) was added to improve exosome production, and exosome was extracted by the ultracentrifugation centrifugation method.^15^ The saucer-like morphology of exosomes was visualized using transmission electron microscopic (TEM) imaging, and the presence of characteristic exosome markers, such as tumor susceptibility gene 101 (TSG101), CD9, and CD81 confirmed the successful isolation of exosomes by western blotting (Figure 1B-C).^16^ Drawing upon previous findings, we postulated that high fatty acid treated adipocyte-derived exosomes (ad-exos) would exhibit enhanced uptake by liver cancer cells.^17^ To corroborate this hypothesis, we co-cultured two distinct types of ad-exos with Hepa1-6 cells. Compared to untreated ad-exos (termed NEM; Normal adipocyte Exosome Membrane), high fatty acid treated ad-exos (termed HEM; High fatty acid treated adipocyte Exosome Membrane) demonstrated increased internalization by HCC cells (Figure 1D-E). Based on the experimental results presented above, it is evident that “obese” ad-exos are significantly more readily absorbed by HCC cells when compared to normal ad-exos. This finding serves as a foundational basis for our subsequent experimental designs.

**Figure 1.**
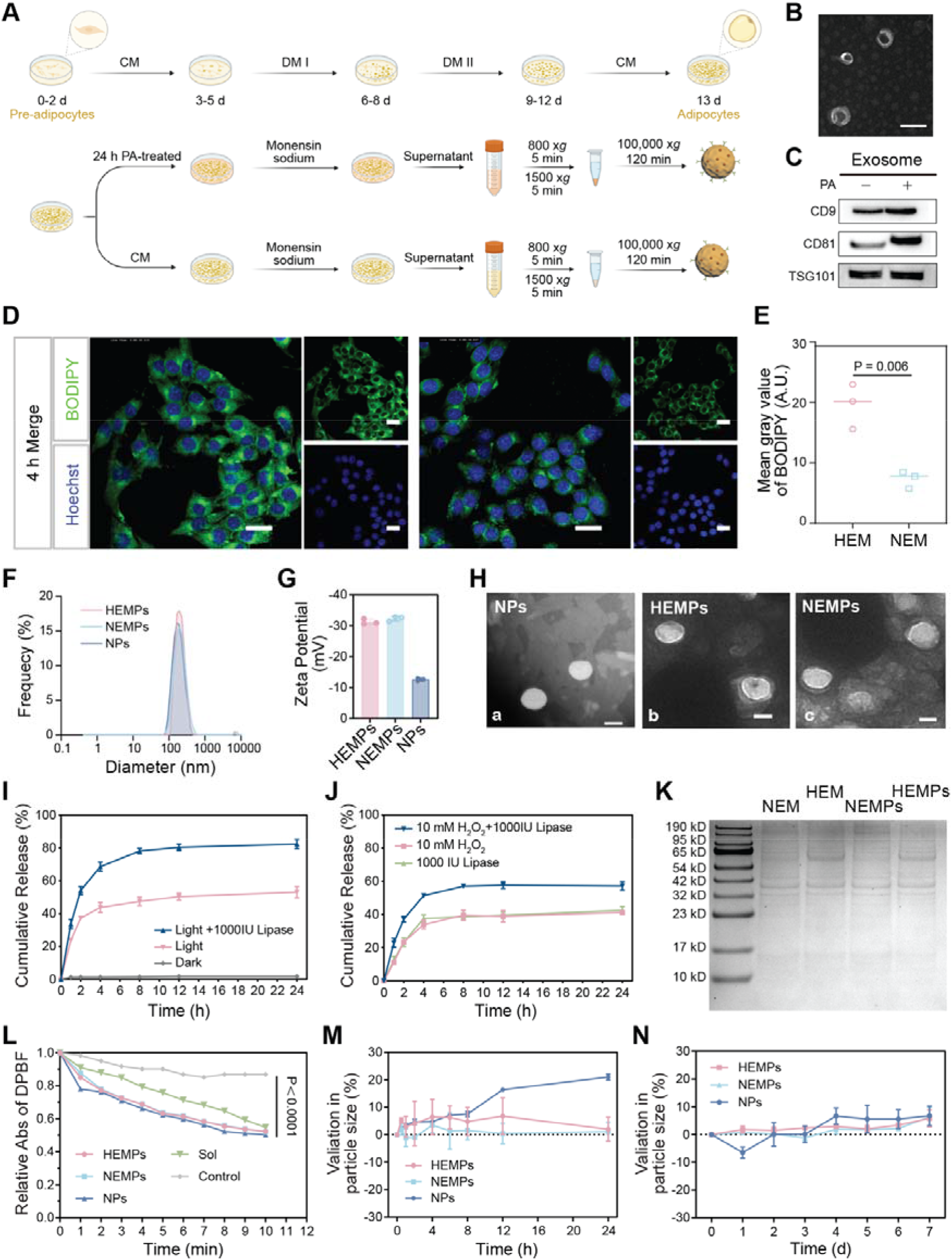
Establishment of hybrid adipocyte-derived exosome nano platform. (A) Schematic illustration of the “cocktail” method to induce differentiation of 3T3-L1 mouse preadipocytes into mature adipocytes, and obtain exosomes secreted from mature adipocytes. (B) A micrograph of transmission electron microscopy (TEM) of exosomes released from 3T3-L1 adipocytes. Scale bar represents 100 nm. (C) Western blotting to evaluate the expression of exosome markers TSG101, CD9, and CD81 in 3T3-L1 adipose-derived exosomes. (D, E) Fluorescent micrographs showing cellular uptake of Hepa1-6 cells treated with HEM (High fatty acid treated adipocyte Exosome Membrane) and NEM (Normal adipocyte Exosome Membrane) at 4 h (D), followed by quantitative image analysis (E). Scale bars represent 20 μm. (F) Histograms to show the hydrodynamic sizes of NPs, HEMPs and NEMPs. (G) A bar graph to show zeta potentials of NPs, HEMPs and NEMPs. (H) Micrographs of transmission electron microscopy of NPs, HEMPs and NEMPs. Scale bars represent 100 nm. (I, J) Cumulative DTX released from lipid core nanoparticles treated with light and 1000 IU lipase (I) in combination with 10 mM H_2_O_2_ (J). (K) SDS-PAGE analysis of protein expressions in HEM, NEM, HEMPs and NEMPs. Left lane shows protein markers. (L) *In vitro* ^1^O_2_ production from NPs, HEMPs, NEMPs and solution (Sol), determined by 1,3-diphenylisobenzofuran (DPBF). (M, N) Evaluation of colloidal stabilities (M) and stabilities (N) of NPs, HEMPs and NEMPs by measuring particle sizes. Data represent a mean ± SD (*n* = 3).

### Design, Preparation and Characterization of the Lipid Core Nanoparticles

To further improve the therapeutic effect of “obese” ad-exos, we designed lipid core nanoparticles to substitute the nuclear neutral lipids in these exosomes, based on the structural characteristics of “obese” ad-exos (Figure S2-S5). As shown in Figure S6A-B, no significant change was found in the peak position and strength of both the UV-Vis and fluorescence spectra, suggesting marginal effect of chemical modifications on the optical properties of PpIX.

Elevated concentrations of ROS and lipases within tumor cells, compared with normal cells, foster an intracellular microenvironment by enhanced oxidation and pronounced lipolysis.^18^ Drawing the distinct difference of normal and tumor cells, a lipid core prodrug was synthesized with oxidation-sensitive thioether bond and lipase-sensitive triglyceride backbone, expected to improve the specific release of prodrug in tumor cells. We further examined the *in vitro* drug release kinetics of lipid core nanoparticles. As shown in Figure 1J, in the presence of 10 mM H_2_O_2_ (a prevailing analogue of ROS) or 1000 IU lipase only, a small amount of DTX release (<50%) from prodrug nanoparticles within 8 h. However, in the presence of both lipase and H_2_O_2_, there was a substantial augmentation in DTX release compared with the release under single trigger conditions. The DSTG prodrug exhibits significant hydrophobic characteristics. The inherent hydrophobicity of these structures and the stability of the nanoparticles serve to protect the thioether bond from oxidative degradation by H_2_O_2_, thus allowing for a controlled release of DTX release. Consequently, lipid core nanoparticles demonstrated a preferential enhancement of drug release under dual trigger condition involving oxidation and lipase, in contrast to a more limited release rate observed under single trigger conditions. According to the release results, we summarized the programmable drug delivery mechanism of DSTG prodrug (Figure S7). When the ester bond in *sn-1* was hydrolyzed in the presence of lipase, increasing the hydrophilicity and disrupting the spatial packing of the resultant structure, which in turn facilitates the subsequent ROS-mediated attack. Moreover, the combined application of lipase and light irradiation induce self-facilitated activation of prodrug, thereby efficiently releasing the DTX drug. Notably, self-amplified synergistic effect on drug release was evident. More than 80% of DTX were released from nanoparticles with light irradiation, which was 1.99 times faster than that in 10 mM H_2_O_2_ medium without light irradiation (Figure 1I). These results confirmed our hypothesis that self-amplified drug release was achieved by self-sacrificed single thioether bond and light stimulation.

### Hybrid Adipocyte-Derived Exosome Nano Platform Promote Tumor Cell Death

To further enhance targeting ability and biocompatibility of the nano platform, we integrated lipid core nanoparticles with ad-exos membrane camouflage. First, the one-step nanoprecipitation method was applied to prepare the lipid core nanoparticles.^19^ Then, the lipid core nanoparticles were extruded with the equal quality of high fatty acid treated ad-exos membrane or normal ad-exos membrane to prepare HEMPs and NEMPs (HEM and NEM cooperated nanoparticles, respectively). As shown in Figure 1F, the average diameters of HEMPs (183.8 ± 3.3 nm; PDI: 0.176) and NEMPs (181.1 ± 4.9 nm: PDI: 0.141) were about 50-nm larger than that of the lipid core nanoparticles (132.5 ± 1.1 nm; PDI: 0.061). The zeta potential of HEMPs (31.2 ± 0.9 mV) and NEMPs (32.3 ± 0.8 mV) was lower than the lipid core NPs (12.6 ± 0.5 mV) (Figure 1G). The increased hydrodynamic diameter and decreased zeta potential indicated the lipid core nanoparticles were successfully covered with the HEM and NEM. The TEM images clearly revealed the core-shell structure of HEMPs and NEMPs, whicle NPs display a bare spherical morphology, further indicating the successful camouflage by the ad-exos membrane (Figure 1H). Moreover, the unique proteins inherited from exosome membranes were well-preserved in the protein profile of ad-exo nanoplatform, indicating that the preparation of HEMPs and NEMPs didn’t affect the key exosome membrane proteins, confirmed by SDS-PAGE (Figure 1K). The self-assembled nano-prodrugs in ad-exos membrane possessed excellent storage stability in PBS at 4°C for seven days (Figure 1N) and desirable colloidal stability in PBS supplemented with 10% fetal bovine serum (FBS) for 24 h (Figure 1M).

To quantify the photosensitizer initiate the production of singlet oxygen (^1^O_2_) within our hybrid adipocyte-derived exosome platform upon exposure to light, we used 1,3-diphenylisobenzofuran (DPBF), a ^1^O_2_ marker for its UV absorption irreversibly decreases upon exposure to ^1^O_2_ (Figure 1L). Compared to lipid core nanoparticles, PpIX produced 0.08-fold less ^1^O_2_ in aqueous solutions due to the poor aqueous solubility. Notably, both HEMPs and NEMPs exhibited a more gradual ^1^O_2_ production rate, suggesting that the exosome membrane may afford a protective effect against ^1^O_2_ production.

### Caveolae-mediated endocytosis primary enhanced drug intake

We then examined the liver cancer cell targeting uptake effect of hybrid adipocyte-derived exosome nano platform (Figures 2A and S8-9), and found the cellular uptake followed this pattern: HEMPs > NEMPs > NPs. Additionally, we noted an augmentation in cellular uptake efficiency following a 4-h incubation period relative to that at 1 h, suggesting a time-dependent cellular uptake of the formulation. Consequently, we opted to use a 4-h incubation period for our subsequent cell experiments. To investigate the mechanisms of hybrid adipocyte-derived exosome nanoplatform into the endocytic pathways, specific endocytic inhibitors were used, such as chlorpromazine (CPZ) for clathrin-mediated endocytosis, filipin (FLP) for caveolae-mediated endocytosis and amiloride (AMI) for micropinocytosis.^20^ Besides, Hepa1-6 cells were cultured at 4 °C to probe the influence of energy on cellular uptake. Figure 2B-D showed that low temperature (4 °C) led to markedly reduced internalization of these nanoplatforms, denoting an energy-dependent uptake mechanism. Subsequently, upon the treatments with CPZ (10 μM), FLP (5 μg mL^−1^) and AMI (10 μM), a more pronounced inhibitory effect of FLP relative to CPZ, whereas AMI partially attenuated the uptake of the hybrid adipocyte-derived exosome platform, suggesting at potential contributions from macrophage and phagocytic pathways in their internalization.

**Figure 2.**
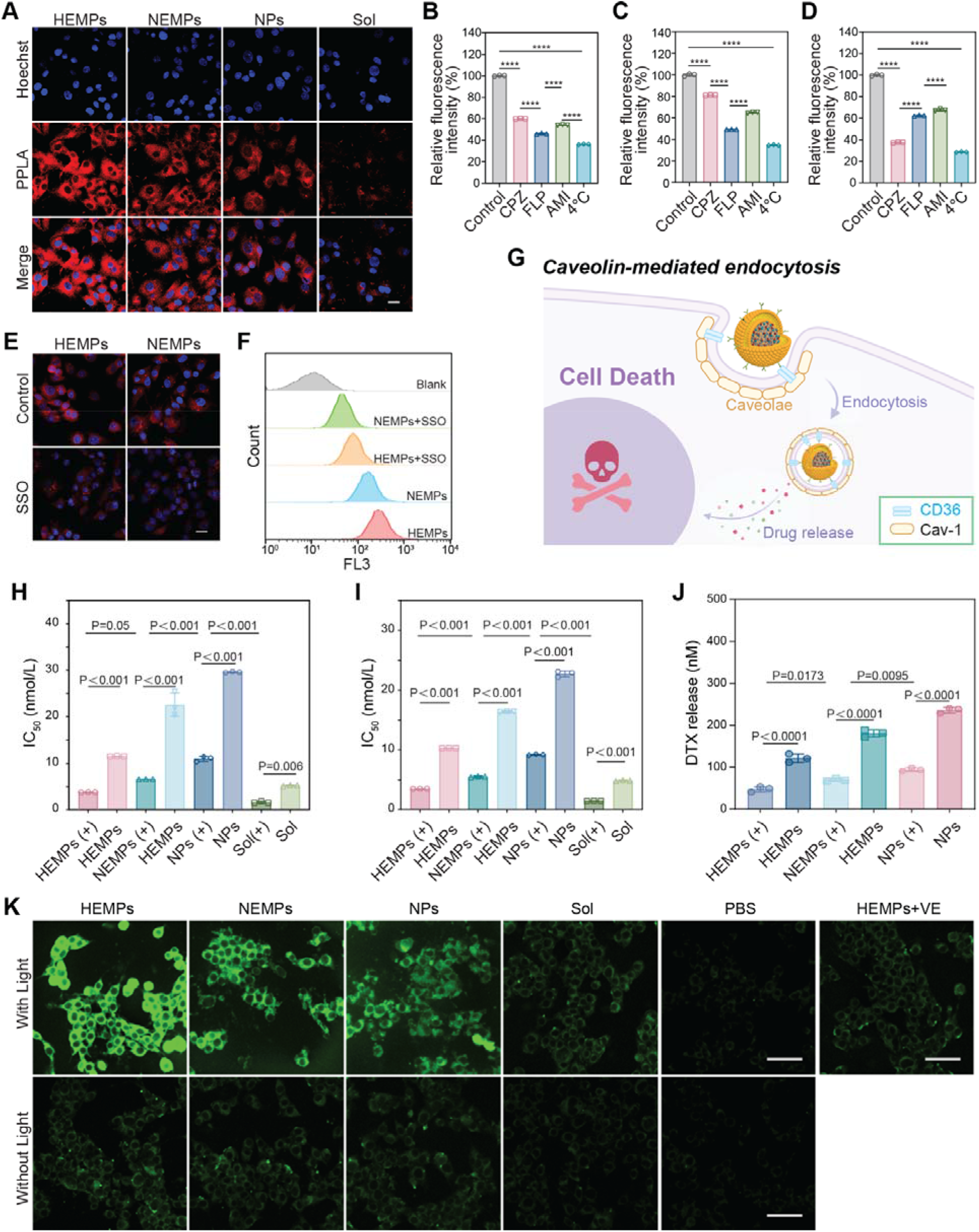
Characterization of a hybrid adipocyte-derived exosome nano platform *in vitro*. (A) Fluorescent micrographs of cellular uptake of mixed solution (Sol), NPs, NEMPs and HEMPs into Hepa1-6 cancer cells at 4 h. Scale bar represents 20 μm. (B, C, D) Flow cytometric analyses of cellular uptake mechanisms of (B) HEMPs, (C) NEMPs, and (D) NPs into Hepa1-6 cells treated with endocytosis inhibitors, chlorpromazine (CPZ), filipin (FLP) and amiloride (AMI). (E) Fluorescent micrographs of cellular uptake in Hepa1-6 cells pretreated with CD36 inhibitor, sulfosuccinimidyl oleate (SSO), followed by introducing HEMPs and NEMPs. Scale bar represents 20 μm. (F) Flow cytometric analysis of cellular uptake of HEMPs and NEMPs in Hepa1-6 cells pretreated with SSO. (G) Schematic illustration of the mechanisms of cellular uptake and drug release via caveolae-mediated endocytosis. (H, I) Evaluation of cytotoxicity of different formulations incubated with (H) HepG2 cells and (I) Hepa1-6 cells with or without light exposure. (J) Evaluation of free DTX released from NPs, NEMPs, and HEMPs after incubation with Hepa1-6 cells. (K) Fluorescent micrographs of ROS produced in Hepa1-6 cells treated with mixed solution (Sol), NPs, NEMPs, and HEMPs. Scale bars represent 50 μm. Data are presented as means ± SD (*n* = 3).

Interestingly, we observed that obese hybrid adipocyte-derived exosome platform, when treated with high fatty acids, exhibited increased uptake by liver cancer cells compared to untreated ad-exos (Figure 2A and S8-9). To investigate this mechanism, we noticed CD36, which was a plasma membrane glycoprotein and worked as a fatty acid transporter,^21, 22^ and confirmed by induced expression of CD36 in ad-exos after high dose of fatty acids (Figure S10). Although efficient cellular uptake through CD36 receptor-mediated transport is uncommon and the mechanism still remains unclear, CD36 could be related to fatty acid uptake. Therefore, we hypothesize that CD36 plays an auxiliary role in the process of cellular endocytosis. To validate this hypothesis, we utilized sulfosuccinimidyl oleate (SSO), a CD36 inhibitor, to evaluate its impact on nanoparticle uptake by HCC cells.^23^ Indeed, SSO pretreatment reduced cellular uptake of hybrid adipocyte-derived exosome (Figure 2E, F). In light of our finding, we summarized the brief caveolae-mediated endocytosis mechanism (Figure 2G).

### In vitro Chemo-Photodynamic Therapy against Cancer Cells

To evaluate the cytotoxicity of hybrid adipocyte-derived exosome platforms on HepG2 human carcinoma cells and Hepa1-6 mouse liver carcinoma cells, we obtained the half maximal inhibitory concentration (IC_50_) values (Figure 2H-I and Table S1). Compared with mixed solution (Sol), prodrug nanoparticles exhibited lower cytotoxicity due to the delayed release of DTX. The cytotoxicity of prodrug nanoparticles followed the other of HEMPs (+)> NEMPs (+)> NPs (+). The same pattern was observed in the non-light-treated group. Besides, the groups without light exposure showed limited cytotoxicity but exhibited effective photodynamic cytotoxicity under light irradiation, indicating the therapy efficacy of PDT. Obviously, the hybrid adipocyte-derived exosome nano platform showed distinct advantage owing to the self-facilitated dual-responsiveness and enhanced cellular uptake effect: (i) self-facilitated and dual responsiveness, the endogenous ROS overproduced in tumor cells and PPLA-generated ROS together with lipase responsiveness synergistically promoted rapid and precise drug release; and (ii) enhanced cellular uptake, the high fatty acid ad-exo nanocarriers improved the cellular uptake and facilitated drug delivery, leading to higher cytotoxicity in nanoprodrugs compared to naked nanoparticles. It has been found that cytotoxicity is closely related to the release of DTX from prodrug nanoparticles: much drug release coincides with higher cytotoxicity. Therefore, the release of DTX from prodrug nanoparticles was evaluated after incubation with Hepa1-6 cells (Figure 2J and S7). Light irradiation generated significant amounts of ROS, promoting the cleavage of single thioether linkage and thereby accelerating the release of DTX. Remarkably, the DTX release from HEMPs was approximately 8.2-fold higher than that from NPs under the same light conditions. These intracellular release results substantiate our initial hypothesis.

### Cellular ROS and Lipid Peroxidation Assessment

Since ROS generation in photodynamic therapy is the most crucial factor for cell destruction, we investigated the production of ROS in various groups of preparations with or without light. Then, we employed the DCFH-DA fluorescent dye to assess the generation of cellular ROS in response to various formulations after a 4 h incubation with Hepa1-6 and HepG2 cells (Figure 2K and S11-12). Cells subjected to light irradiation exhibited notably intensified green fluorescence, whereas those without light exposure displayed minimal fluorescence signals. Notably, HEMPs exhibited a substantially higher ROS production compared to NPs, likely due to the highly efficient uptake facilitated by CD36-aided endocytosis. This quantitative assessment of cellular ROS generation in both Hepa1-6 and HepG2 cells was corroborated using Image J, aligning with the observations made through confocal fluorescence microscopy and inverted microscope imaging. Our research also indicates that vitamin E (VE), as a lipid-soluble ROS scavenger,^13^ can effectively alleviate the changes in intracellular ROS levels caused by our formulations. The phospholipid bilayer structure of ad-exos, along with unsaturated fatty acids enriched in lipid core nanoparticles, serves as fuels for fatty acid oxidation in the presence of high levels of ROS. We assumed that ROS generated by HEMPs could oxidize the massive amounts of lipids inside the cancer cells, resulting in lipid peroxides accumulation, which further promoted the cell death. Consequently, we further explored the impact of lipid peroxidation using the C11 BODIPY 581/591 probe (Figure S13). The obese hybrid adipocyte-derived exosome displayed substantially elevated levels of lipid peroxidation in comparison to the NPs group, which suggests that both ROS and LPO mechanisms would enhance the anti-tumor effects.

### Prolonged Cellular Drug Retention and Enhanced Drug Targeted Tumor Accumulation

Due to the tumor-targeting and self-protective properties of obese hybrid adipocyte-derived exosome, we hypothesized that HEMPs could enhance drug retention in the bloodstream and improve tumor specific distribution. In addition, Figure 1M shows that exosomes of hybrid adipocytes have good colloidal stability, so this provides a good basis for our hypothesis. To investigate this, we examined the pharmacokinetics of Mixed solutions, NPs, HEMPs, and NEMPs in Sprague-Dawley rats (Figure 3B-D and Table S2). Mixed solutions were quickly eliminated from the bloodstream, duo to their short half-life (t_1/2_, 0.18 h), consistent with previous findings.^24^ In contrast, HEMPs and NEMPs extended blood circulation, with intriguing results: the area under the curve (AUC_0–24h_) for prodrugs in NPs, NEMPs, and HEMPs was 18.5-fold, 35.7-fold, and 53.3-fold higher than that of free mixed solutions (Figure 3C). Further, NPs had lower AUCs compared with HEMPs and NEMPs due to their poor colloidal stability (Figure 3A and 1M). This extended circulation time is known to enhance prodrug accumulation in tumors through the enhanced permeability and retention (EPR) effect. The pharmacokinetic results suggested that the obese hybrid adipocyte-derived exosome distinctly prolonged their circulation time in blood.

**Figure 3.**
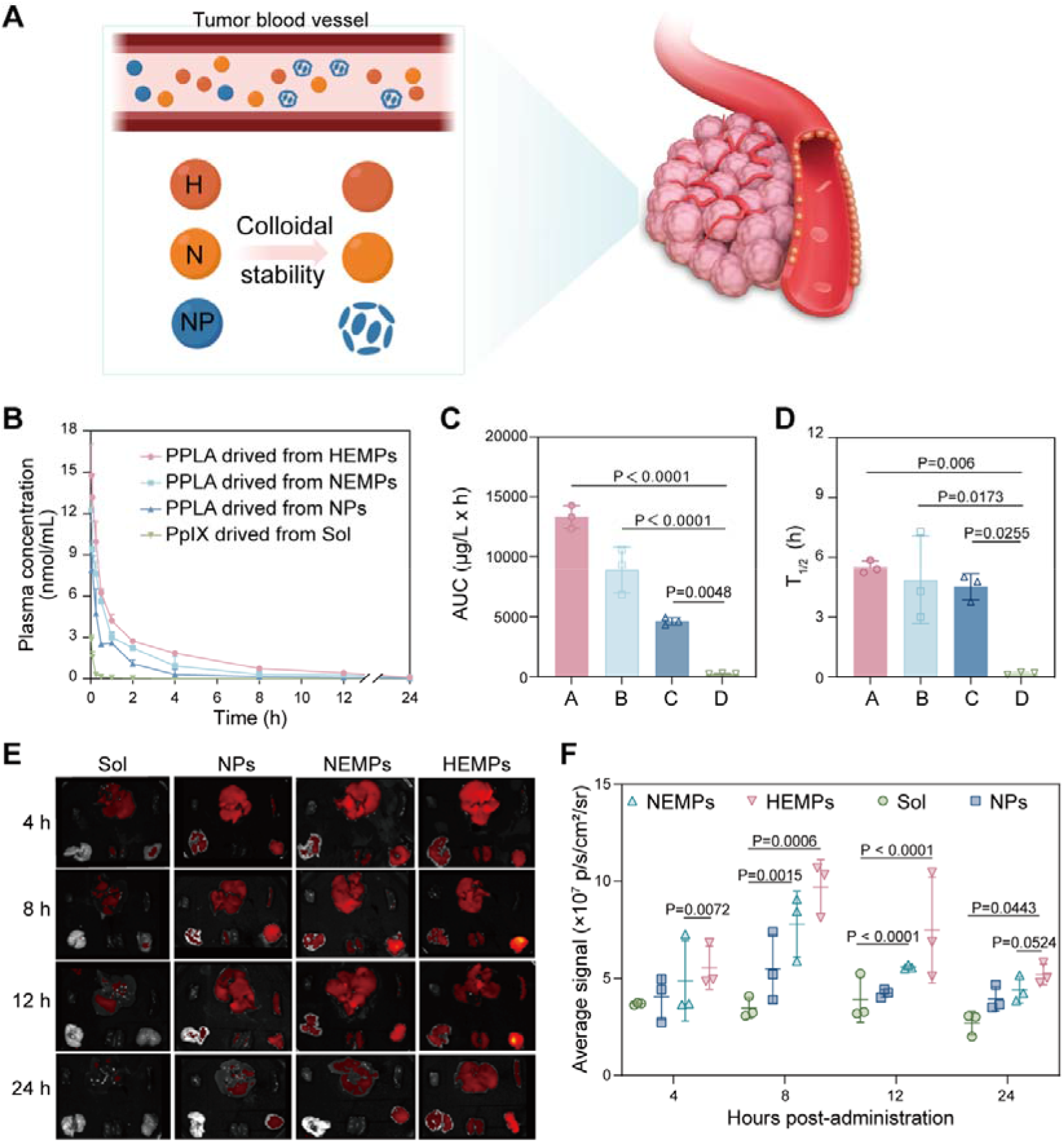
Pharmacokinetics and tumor accumulation of different formulations. (A) Schematic illustration of blood circulation of HEMPs (H), NEMPs (N) and NPs. (B) Molar concentration-time curves of PPLA in rats treated with HEMPs, NEMPs, NPs, and mixed solution (Sol). (C-D) AUC and T_1/2_ parameters of different formulations. (A termed HEMPs, B termed NEMPs, C termed NPs, and D termed Sol) (E) Fluorescence images of PPLA distribution at different time interval. (F) Quantitative analysis of relative organ and tumor accumulation at 4, 8, 12 and 24 h by IVIS spectrum small-animal *in vivo* imaging system. Data are presented as a mean ± SD (*n* = 3).

To achieve effective *in vivo* antitumor results and alleviate drug toxicity, it’s imperative to concentrate nanotherapeutics selectively within tumors while sparing healthy organs. Hence, prior to conducting therapeutic experiments, we investigated the tumor-targeting ability and time-dependent accumulation of hybrid adipocyte-derived exosome in Hepa1-6 tumor-bearing mice (refer to Figure 3E-F). Thanks to the co-assembled NIR fluorescent probe PPLA, we were able to track various nanoparticles *in vivo* without the need for additional fluorescent dye labeling. After injection, we extracted major organs and tumors for *ex vivo* imaging. Notably, the presence of PPLA rapidly diminished in the bloodstream upon administering free PPLA solution, likely due to systemic elimination, resulting in reduced tumor accumulation. As depicted in the Figure 3E-F, the fluorescence intensity of HEMPs increased over time, peaking at 8 hours, notably stronger than that of NPs. This trend was consistent in the quantitative fluorescence results of tumor tissues. Furthermore, the *ex vivo* fluorescence of HEMPs in tumor tissue remained elevated at 24 hours post-administration, indicating a 1.32-fold increase compared to that of NPs. These findings underscore the dual benefits of high fatty acid ad-exos camouflage: not only does it enhance tumor accumulation, but it also augments the accumulation of HEMPs within the tumor.

### *In Vivo* tumor inhibition in Hepa1-6-subcutaneous xenograft model

Encouraged by promising tumor accumulation, we proceeded to investigate the *in vivo* antitumor effectiveness of hybrid adipocyte-derived exosome in Hepa1-6 tumor-bearing mice (Figure 4A). For photodynamic treatment, we subjected the tumors to light irradiation (660 nm, 100 mW cm^−2^) for 2 min, 8 h after administration. Intravenous administration of hybrid adipocyte-derived exosome and subsequent irradiation at 660 nm yielded remarkable tumor growth inhibition, confirmed with changes of body weights (Figure 4B), tumor volumes (Figure 4C), tumor burden (Figure 4D) and actual tumor specimens (Figure 4E). Also, the HEMPs (+) group exhibited excellent tumor inhibition capacity (approximately 210 mm^3^ on day 11). This outcome highlights the advantages of our approach, which combines photodynamic therapy, ROS-mediated enhanced DTX chemotherapy, and the exceptional tumor targeting capability facilitated by obese ad-exo interaction. After light irradiation treatment, all experimental groups exhibited significant tumor growth suppression. Conversely, the free solution group displayed minimal tumor growth suppression in the absence of light irradiation, which may be attributed to the elimination by the reticuloendothelial system. Among the various therapeutic approaches tested, HEMPs combined with light irradiation therapy demonstrated potent antitumor activity. Here, we conclude that this hybrid adipocyte-derived exosome nanoplatform was safe and exerted superior antitumor efficiency because this approach only specifically activated toxic drugs at tumor site after exposure to light irradiation and did not induce systemic toxicity.

**Figure 4.**
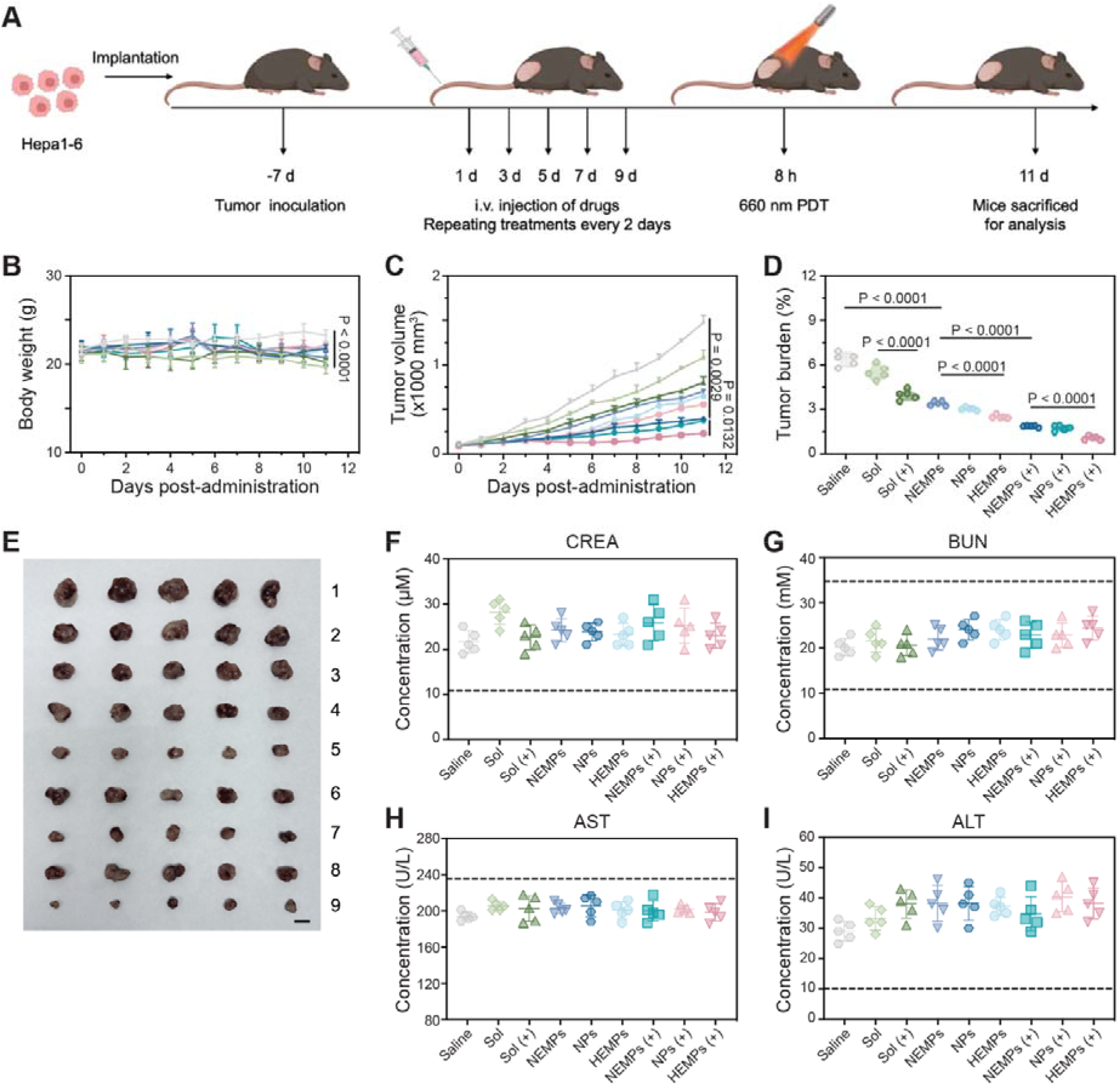
In vivo antitumor activity of HEMPs, NEMPs, NPs, and mixed solution (Sol) in Hepa1-6-subcutaneous xenograft model. (A) Schematic illustration of the tumor treatment against Hepa1-6 xenograft tumor (*n* = 5). (B) The body weight, (C) tumor growth curves, (D) tumor burden, and (E) tumor images of different formulations. Scale bar represents 1 cm. (F-I) Serum biochemical analysis of liver and kidney damaged markers, creatinine (CREA, F), urea nitrogen (BUN, G), aminotransferase (ALT, H) and aspartate aminotransferase (AST, I) (n = 5 mice per group). Data are presented as means ± SD (n = 5).

To evaluate the safety of various formulations, we examined body weight and hepatorenal function parameters. In all formulation groups, there were no discernible alterations in these parameters, indicating excellent biocompatibility, by measuring tissue damage markers, such as creatinine (CREA), urea nitrogen (BUN), aminotransferase (ALT) and aspartate aminotransferase (AST) (Figure 4F-I, respectively). In summary, our hybrid adipocyte-derived exosome nanoplatform has been demonstrated to be both safe and highly effective in targeting tumors. This innovative approach selectively activates toxic drugs at tumor sites following light exposure, without causing any systemic toxicity. Benefit from the advantages of targeted delivery, enhanced circulation, and biocompatible properties offered by exosomes. Our self-facilitated drug release process, coupled with multimodal therapy, guarantees both effectiveness and safety.

## Discussion

Multi-modal cancer therapy leveraging nanodrug delivery systems presents a suite of advantages over conventional monotherapies, notably the potential for synergistic antitumor effects. This is particularly pertinent in the treatment of HCC, where traditional therapies often fall short, underscoring the critical need for innovative therapeutic strategies. Despite its promise, this approach grapples with challenges such as the variability in pharmacokinetics and biodistribution of the drugs involved, circulatory concerns, and the potential for unintended effects on healthy tissues. To surmount these hurdles, we have devised a strategy employing a hybrid nano platform derived from adipocyte exosomes. This platform harnesses dual nano prodrugs for chemotherapeutic and phototherapeutic action, encapsulated within a lipid nano core, which is then integrated into an adipose-derived exosome to enhance tumor targeting, enable dual-responsive and self-amplifying drug release mechanisms, and improve *in vivo* stability. Consequently, our approach has yielded significantly improved therapeutic outcomes, exhibiting commendable biocompatibility and biodegradability under physiological conditions, and demonstrating robust antitumor activity.

In implementing this strategy for HCC treatment, the selection of the exosome source is paramount. Recent literature has highlighted the therapeutic potential of exosomes not only in cancer treatments but also in managing other complex diseases^25^ but have also shown promise in treating other complex diseases.^26^ In our study, we opted for adipocyte-derived exosomes, aligning with findings that link obesity to an increased risk of HCC.^11^ By carefully extracting the exosome from adipocytes,^19^ we secured membranes with preserved expression of proteins characteristic of high fatty acid treatment (Figure 1), which facilitated targeted drug delivery via CD36-auxiliaried clathrin-mediated endocytosis (Figure 2) —a pathway implicated in macrophage and phagocytic activity and known to enhance the uptake of exosomes by HCC cells, thereby promoting tumor progression.^21, 27^

However, the integration of drugs into exosomes without compromising their integrity remains a challenge. Our solution involves self-assembled prodrug nanosystems, which offer enhanced pharmacokinetics and precise control over drug release.^5^ Aforementioned, we utilized “obese” ad-exos with a lipid-rich milieu to incorporate hydrophobic molecules like PUFAs and TGs as the building blocks of our nano prodrugs. This strategy mitigated concerns regarding the disruption of the exosomal structure post drug incorporation.

Furthermore, we explored the use of ROS as a trigger for drug release, exploiting the transient nature of ROS in physiological conditions to achieve spatial and temporal precision to regulate cellular functions.^28^ Specifically, we utilized the short-lived ^1^O_2_ with a lifetime of 3 µs in cultured cells to control drug release through ROS-activated linkages. Our findings confirmed the release of ^1^O_2_ post light exposure, which was absent without light or in the absence of nanodrugs (Figure 2K), illustrating a “self-amplifying cascade” facilitated by the ROS-rich tumor microenvironment.

While our results underscore the efficacy of the hybrid adipocyte-derived exosome nano platform *in vivo*, several challenges persist. First, the replacement of the original exosomal core with nano prodrugs eliminates the inherent cellular components like proteins, RNA/DNA and lipids,^29^ which could otherwise reprogram cellular functions. For instance, exosomes from M1 macrophages can bolster antitumor immunity,^30^ a benefit we forfeit with our current design. Moreover, the complexity of the components and procedures involved poses a challenge for clinical predictability. Simplifying this complexity remains a priority to translate this promising approach into a viable multi-modal cancer therapy.

## Conclusion

In summary, by exploiting an approach utilizing adipocyte-derived exosome derivatization and prodrug nanoparticles, we have successfully developed hybrid adipocyte-derived exosome nanoplatform from therapeutic prodrug cocktails that are remotely activated by NIR light and achieve synergistic chemo-photodynamic therapy to induce cancer remission. The hybrid adipocyte-derived exosome nanoplatform includes several features: (i) Structural Advantage: Utilizing triglyceride’s structural backbone and hydrophobic side chains, we’ve created a lipid-like core structure with assembly capacity for prodrugs. (ii) Release Advantage: This unique design renders the nanocarrier dual-responsive to both lipase and reactive oxygen species. Upon near-infrared (NIR) photoirradiation, the photosensitizer generates ROS, which spontaneously facilitates the neighboring self-immolation of single thioether bond to activate the cytotoxic drug docetaxel, thereby accelerating tumor cell death. (iii) Multimodal Therapeutic Efficacy: The synergistic convergence of DTX-initiated chemotherapy, PpIX-mediated photodynamic therapy, and lipid peroxidation effects expedites the progress of cancer treatment. (iv) Targeting Advantage: Our research findings indicated that the anti-tumor effect of the obese hybrid adipocyte-derived exosome nanoplatform exhibited significantly potent antitumor effect compared to the untreated counterpart. This enhancement could be attributed to the CD36-auxiliaried caveolae-mediated endocytosis. However, the specific mechanism underlying this phenomenon remains unclear and requires further investigation.

As a result, DTX-initiated chemotherapy combining PpIX-mediated PDT demonstrated high-efficiency multi-model antitumor therapy in hepatocellular carcinoma xenograft model. The hybrid adipocyte-derived exosome nanoplatform, integrating multiple drug delivery technologies into one platform with high-efficient targeting and co-delivery characteristics, provides a new strategy for high-efficient multimodal cancer therapy.

## Experimental Section

### Materials

Protoporphyrin IX (PpIX) was purchased from Shanghai Xianhui Pharmaceutical Co., Ltd. (Shanghai, China). Docetaxel was obtained from Dalian Meilun Biotechnology Co., Ltd. (Dalian, China). Linoleic acid was purchased from Shanghai Bide Pharmatech Ltd. (Shanghai, China). The anti-CD63 antibody (A19023), anti-CD36 antibody (A19016), anti-CD9 antibody (A19027), anti-GAPDH antibody (A19056) were purchased from Abclonal Biotechnology Co., Ltd. (Wuhan, China). The anti-CD81 antibody (ab109201) was obtained from Abcam Biotechnology Co., Ltd. (Shanghai, China). DMEM cell culture medium (MA0212) was purchased from Dalian Meilun Biotechnology Co., Ltd. (Dalian, China). Fetal Bovine Serum (PWL001) was purchased from Dalian Meilun Biotechnology Co., Ltd. (Dalian, China). ECL (MA0186) was purchased from Dalian Meilun Biotechnology Co., Ltd. (Dalian, China). DEPE-PEG was purchased from AVT (Shanghai) Pharmaceutical Tech Co., Ltd. Sulfosuccinimidyll Oleate (SSO, a CD36 inhibitor) was obtained from Shanghai Weihuan Biotechnology Co., Ltd. (Shanghai, China).

### Synthesis of PPLA and DSTG

Briefly, we synthesized PPLA through two esterification reactions (see Figure S2) and DSTG via a four-step process (see Figure S3). The final prodrugs were validated using ^1^H NMR and mass spectrometry (see Figure S4-5).

### Ultraviolet and fluorescence spectra

PPIX and PPLA (2 μg mL^−1^, PPIX equivalent) dissolved in dimethylsulphoxide (DMSO) and their ultraviolet were scanned by varioskan lux multimode microplate reader and the fluorescence spectra was measured at the excitation wavelength of 408 nm and emission wavelength of 630 nm.

### Photodynamic efficiency

DPBF in DMSO (6 μL, 5 mM) was added to aqueous solutions (3 mL) of free PPIX, PDSG NPs (20 µg mL^−1^ equivalent PPIX). The absorption spectrum of the mixtures under 660 nm light irradiation (100 mW cm^−2^) was obtained on a varioskan lux multimode microplate reader every 1 min.

### Cell culture and induction of adipocyte differentiation

The mouse preadipocyte cell line 3T3-L1, HepG2 cells and Hepa1-6 cells were cultured in DMEM with 10% (*v*/*v*) fetal bovine serum (FBS) and 1% (*v*/*v*) penicillin-streptomycin at 37 °C under a humidified atmosphere containing 5% CO2. After reaching confluence on day 0, we induced differentiation of 3T3-L1 cells using DMEM with 10% (*v*/*v*) FBS and DMI (IBMX 0.5 mM, DEX 1 μM, Rosiglitazone 2 μM, and insulin 5μg mL^−1^) for 2 days. Subsequently, we replaced the culture medium with DMEM containing 10% FBS and DMII (insulin 5 μg mL^−1^) for another 2 days. This was followed by a 4-day incubation in DMEM with 10% (*v*/*v*) FBS, with medium changes every 2 days. Mature adipocytes emerged by day 12 and were identified by their red staining with Oil Red O. The cells were fixed with 4% (*v*/*v*) paraformaldehyde for 15 min, followed by a rinse with PBS. They were then treated with 60% (*v*/*v*) isopropanol for 5 min, stained with Oil Red O for 1 h, and washed four times with distilled water. Finally, the stained cultures were photographed at 40× magnification using an inverted fluorescence microscope.

### Preparation of HMNPs and NMNPs

The PPLA and DSTG co-assembled nanoparticles were prepared by the simple one-step nanoprecipitation method. Adipocyte-derived exosomes were collected by culturing adipocytes in MON medium for 4 h. Afterward, we removed cells with a 300 ×*g* centrifugation for 10 min and cleared cell debris with a 15,000 ×*g* centrifugation for 30 min. Further ultracentrifugation at 100,000 ×*g* for 120 min yielded exosome pellets. These pellets were resuspended in a hypotonic buffer with a protease inhibitor cocktail and stored at 4 °C for 12 h. The mixture underwent a 100,000 ×*g* ultracentrifugation for 2 h to obtain an exosome membrane. For the creation of exosome biomimetic HEMPs, we repeatedly coextruded a mixture of nanoparticles and exosome membranes using a 220 nm polycarbonate porous membrane on an extruder from Avestin Inc., Ottawa, Canada. The same procedures were applied to fabricate NEMPs.

### In vitro ROS generation

The DPBF was dispersed in different samples at a PPIX dose of 20 μg mL^−1^. When exposed to 660nm light irradiation (100 mW cm^−2^), the ultraviolet absorption in 410 nm was detected every 60 s to monitor singlet oxygen level.

### In vivo ROS generation

Intracellular ROS production was detected with DCFH-DA staining by inverted fluorescence microscope imaging, Hepa1-6 cells and HepG2 cells (80,000 cells/well) were cultivated in 24-well plates for 24 h. HEMPs, NEMPs, NPs and Sol were added to cells for 4 h. Subsequently, cells were stained with DCFH-DA for 0.5 h and washed with PBS 3 times. The DCF fluorescence was detected with inverted fluorescence microscope and confocal fluorescence microscope.

### Cytotoxicity assay

Cell cytotoxicity was evaluated using MTT assay. Briefly, Hepa1-6 and HepG2 cancer cells were seeded in 96-well plates (2,000 cells/well) for 12 h and treated with free mixed solution, NPs, HEMPs, and NEMPs with a series of concentrations for 4 h. Subsequently, the cells were exposed to a laser (660 nm, 50 mW cm^−2^) for 2 min or incubated for another 44 h. Subsequently, 20 μL of MTT (5 mg mL^−1^) solution was added and then incubated at 37 °C. After 4 h of incubation, the formazan crystals were dissolved in 200 μL of DMSO. Finally, the absorbance of samples was evaluated using Varioskan Flash multimode microreader.

### Lipid peroxidation evaluation

To evaluation of intracellular lipid peroxidation levels of treated with different formulations, the BODIPY 581/591 C11 fluorescence probe was used. Hepa1-6 cells were treated with different formulations for 4 h. After that, the media was substituted with fresh medium containing BODIPY probe (5 μM) for 30 min. Finally, the excess probe was discarded, and the cells cleaned with PBS before observed by confocal microscopy.

### Flow cytometry

To conduct cellular uptake, Hepa1-6 and HepG2 cells with 80%-90% confluence was cultured with fresh medium containing different formulations for 1 and 4 h. After that, the cells washed with PBS, and harvested after trypsinization. The collected cells were centrifugated and resuspended in PBS buffer to remove medium and trypsin. Flow cytometry analyses were subsequently carried out on FL3-H flow cytometer.

### Western blotting

Radio Immunoprecipitation Assay (RIPA)-lysis buffer was used to acquire exosome lysates. The proteins were fractionated with SDS-PAGE electrophoresis, transferred to PVDF and incubated with primary antibodies: CD36, CD63, CD9, CD81, and GAPDH at 4 LJ overnight. Following secondary antibody incubation, ECL Western Blotting Substrate was added wo visualize the protein bands.

### In vitro drug release

The release behavior of DTX from PDSG NPs was evaluated by HPLC method. To test the ROS sensitivity, hydrogen peroxide (H_2_O_2_) was applied as an ROS simulant. Lipase was added to realistically mimic the tumor microenvironment. With PBS (pH 7.4) containing 30% (*v*/*v*) ethanol as the release medium, the release of docetaxel from lipid drop-like prodrug nanoparticles was investigated with or without H_2_O_2_ (10 mM) or lipase (1000IU mL^−1^), and with light (100mW cm^−2^, 2 min) or light avoidance. At different time points, 1 mL of dialysis was withdrawn to investigate the release behaviors of DTX using HPLC.

### Animal studies

All experiments were performed in accordance with the guide approved by the Institutional Animal Ethical Care Committee (IAEC) of Shenyang Pharmaceutical University.

### Pharmacokinetics and Biodistribution

The pharmacokinetic behaviors of free mixed solution, PDSG NPs, NEMPs and HEMPs were determined using Sprague-Dawley rats (200-230 g). The Sprague-Dawley rats were divided into 4 groups (three rats per group). Formulations were injected into the tail vein with an equivalent dose at 2 mg kg^−1^ PpIX and 2 mg kg^−1^ DTX. At designed time points, about 500 μL of blood was collected and centrifuged at 15,000 rpm and 150 μL of plasma was obtained. The concentration of PPLA was analyzed using Varioskan Flash multimode microreader. The biodistribution and tumor accumulation of formulations was detected on Hepa1-6 tumor-bearing C57BL/6 mice. At the tumor volume up to 300 mm^3^, free mixed solution, PPLA-DSTG NPs, NEMPs and HEMPs were intravenously administrated with the equivalent dose at 2 mg kg^−1^ PpIX and 2 mg kg^−1^ DTX (3 mice per group). After 4, 8, 12 and 24 h post injection, the fluorescence signals in the mice were detected using the Spectrum In Vivo Imaging System (IVIS).

### In vivo antitumor efficacy

Mice bearing Hepa1-6 tumor were subcutaneously injected with 100 μL of Hepa1-6 cells (5_×_10^6^ cells per mouse) on the right limbs. The tumor-bearing mice were divided into 9 groups (5 mice per group). When the tumor volume reached about 100 mm^3^, formulations were intravenously administrated into mice with an equivalent dose at 2 mg kg^−1^ PpIX and 2 mg kg^−1^ DTX. For photodynamic treatment, the tumors were treated by NIR light irradiation (660 nm, 100mW cm^−2^) for 5 min after administration 8h. The tumor volume was defined as *V* = *W*^2^*L*/2, where the *W* and *L* are the shorted and longest diameters respectively. The relative volume was calculated as *V*/*V*_0_, where *V*_0_ and *V* are the tumor volumes before and after administration, respectively.

### Data availability

All relevant data are available from the authors.

### Declaration of competing interest

The authors declare that they have no known competing financial interests or personal relationships that could have appeared to influence the work reported in this paper.

## Supporting information

Synthesis pathways of PPLA.

Synthesis pathways of DSTG.

(A) Mass spectrum and (B) 1H NMR spectrum in CDCl3 of PPLA.

(A) Mass spectrum and (B) 1H NMR spectrum in CDCl3 of DSTG.

(A)The fluorescence spectra and (B) ultraviolet spectra of PpIX and PPLA.

The oxidative release mechanism of the prodrug nanoparticles.

Cellular uptake of HepG2 cancer cells.

Fluorescence distribution from flow cytometry of Hepa1-6 cells incubated for (A) 1 h or (B) 4 h.

Western blotting of characteristic markers CD36, CD9, CD81, TSG101, and GAPDH. (a: normal treatment; b: palmitic acid treatment)

In vitro cellular lipid peroxidation level.

In vitro cytotoxicity (IC50 values).

Pharmacokinetic parameters of Mixed solution, HEMPs, NEMPs, and NPs.

## Acknowledgements

Funding was generously provided by the National Natural Science Foundation of China (82104109), Liaoning Provincial Department of Education Program (LJKMZ20221353 and LJKZ0940), Natural Science Foundation of Liaoning Province (2022-BS-158) and the Japan Society for the Promotion of Science (JSPS; 21H01728). The WPI-iCeMS is supported by the World Premier International Research Centre Initiative (WPI), MEXT, Japan.

## Author contributions

X.L. conceived the project, wrote the manuscript, J.Z., M.L., Q.X., and B.S. collected citations and drew the scheme, C.T., K.K. revised the manuscript.

## Notes

### Competing Interest Statement

The authors have declared no competing interest.

### Summary of Updates

The modified version adds additional materials and methods sections, remaining the same as the original version.

## Reference

(1) Vogel, A.; Meyer, T.; Sapisochin, G.; Salem, R.; Saborowski, A. Hepatocellular carcinoma. Lancet 2022, 400 (10360), 1345–1362. DOI: 10.1016/S0140-6736(22)01200-4 PubMed. Schaue, D.; McBride, W. H. Opportunities and challenges of radiotherapy for treating cancer. Nat Rev Clin Oncol 2015, 12 (9), 527–540. DOI: 10.1038/nrclinonc.2015.120 PubMed.

(2) Usuda, J.; Kato, H.; Okunaka, T.; Furukawa, K.; Tsutsui, H.; Yamada, K.; Suga, Y.; Honda, H.; Nagatsuka, Y.; Ohira, T.;, et al. Photodynamic therapy (PDT) for lung cancers. J Thorac Oncol 2006, 1 (5), 489–493. PubMed. Zhao, J.; Duan, L.; Wang, A.; Fei, J.; Li, J. Insight into the efficiency of oxygen introduced photodynamic therapy (PDT) and deep PDT against cancers with various assembled nanocarriers. Wiley Interdiscip Rev Nanomed Nanobiotechnol 2020, 12 (1), e1583. DOI: 10.1002/wnan.1583 PubMed.

(3) Ho, W. J.; Jaffee, E. M.; Zheng, L. The tumour microenvironment in pancreatic cancer - clinical challenges and opportunities. Nat Rev Clin Oncol 2020, 17 (9), 527–540. DOI: 10.1038/s41571-020-0363-5 PubMed.

(4) Choi, J.; Sun, I.-C.; Sook Hwang, H.; Yeol Yoon, H.; Kim, K. Light-triggered photodynamic nanomedicines for overcoming localized therapeutic efficacy in cancer treatment. Advanced Drug Delivery Reviews 2022, 186. DOI: 10.1016/j.addr.2022.114344. Lucky, S. S.; Soo, K. C.; Zhang, Y. Nanoparticles in Photodynamic Therapy. Chemical Reviews 2015, 115 (4), 1990–2042. DOI: 10.1021/cr5004198. Li, W.-P.; Yen, C.-J.; Wu, B.-S.; Wong, T.-W. Recent Advances in Photodynamic Therapy for Deep-Seated Tumors with the Aid of Nanomedicine. Biomedicines 2021, 9 (1). DOI: 10.3390/biomedicines9010069.

(5) Ma, S.; Kim, J. H.; Chen, W.; Li, L.; Lee, J.; Xue, J.; Liu, Y.; Chen, G.; Tang, B.; Tao, W.;, et al. Cancer Cell-Specific Fluorescent Prodrug Delivery Platforms. *Advanced Science (Weinheim, Baden-Wurttemberg*, Germany) 2023, 10 (16), e2207768. DOI: 10.1002/advs.202207768 PubMed. Krishnan, N.; Fang, R. H.; Zhang, L. Engineering of stimuli-responsive self-assembled biomimetic nanoparticles. Advanced Drug Delivery Reviews 2021, 179, 114006. DOI: 10.1016/j.addr.2021.114006 PubMed. Zhou, Y.; Li, Q.; Wu, Y.; Li, X.; Zhou, Y.; Wang, Z.; Liang, H.; Ding, F.; Hong, S.; Steinmetz, N. F.; et al. Molecularly Stimuli-Responsive Self-Assembled Peptide Nanoparticles for Targeted Imaging and Therapy. ACS Nano 2023, 17 (9), 8004–8025. DOI: 10.1021/acsnano.3c01452 PubMed. Pei, Q.; Jiang, B.; Hao, D.; Xie, Z. Self-assembled nanoformulations of paclitaxel for enhanced cancer theranostics. Acta Pharmaceutica Sinica. B 2023, 13 (8), 3252–3276. DOI: 10.1016/j.apsb.2023.02.021 PubMed. Deng, Z.; Liu, S. Controlled drug delivery with nanoassemblies of redox-responsive prodrug and polyprodrug amphiphiles. Journal of Controlled Release : Official Journal of the Controlled Release Society 2020, 326, 276–296. DOI: 10.1016/j.jconrel.2020.07.010 PubMed.

(6) Mu, J.; Zhong, H.; Zou, H.; Liu, T.; Yu, N.; Zhang, X.; Xu, Z.; Chen, Z.; Guo, S. Acid-sensitive PEGylated paclitaxel prodrug nanoparticles for cancer therapy: Effect of PEG length on antitumor efficacy. J Control Release 2020, 326, 265–275. DOI: 10.1016/j.jconrel.2020.07.022 PubMed.

(7) Luo, C.; Sun, J.; Liu, D.; Sun, B.; Miao, L.; Musetti, S.; Li, J.; Han, X.; Du, Y.; Li, L.;, et al. Self-Assembled Redox Dual-Responsive Prodrug-Nanosystem Formed by Single Thioether-Bridged Paclitaxel-Fatty Acid Conjugate for Cancer Chemotherapy. Nano Letters 2016, 16 (9), 5401–5408. DOI: 10.1021/acs.nanolett.6b01632 PubMed.

(8) Wang, S.; Wang, Z.; Yu, G.; Zhou, Z.; Jacobson, O.; Liu, Y.; Ma, Y.; Zhang, F.; Chen, Z.-Y.; Chen, X. Tumor-Specific Drug Release and Reactive Oxygen Species Generation for Cancer Chemo/Chemodynamic Combination Therapy. Adv Sci (Weinh) 2019, 6 (5), 1801986. DOI: 10.1002/advs.201801986 PubMed. Ye, M.; Han, Y.; Tang, J.; Piao, Y.; Liu, X.; Zhou, Z.; Gao, J.; Rao, J.; Shen, Y. A Tumor-Specific Cascade Amplification Drug Release Nanoparticle for Overcoming Multidrug Resistance in Cancers. Advanced Materials (Deerfield Beach, Fla.) 2017, 29 (38). DOI: 10.1002/adma.201702342 PubMed. Zhao, X.; Amevor, F. K.; Xue, X.; Wang, C.; Cui, Z.; Dai, S.; Peng, C.; Li, Y. Remodeling the hepatic fibrotic microenvironment with emerging nanotherapeutics: a comprehensive review. J Nanobiotechnology 2023, 21 (1), 121. DOI: 10.1186/s12951-023-01876-5 PubMed.

(9) Flaherty, S. E.; Grijalva, A.; Xu, X.; Ables, E.; Nomani, A.; Ferrante, A. W. A lipase-independent pathway of lipid release and immune modulation by adipocytes. Science 2019, 363 (6430), 989–993. DOI: 10.1126/science.aaw2586 PubMed. Wang, G.; Li, J.; Bojmar, L.; Chen, H.; Li, Z.; Tobias, G. C.; Hu, M.; Homan, E. A.; Lucotti, S.; Zhao, F.; et al. Tumour extracellular vesicles and particles induce liver metabolic dysfunction. Nature 2023, 618 (7964), 374–382. DOI: 10.1038/s41586–023-06114–4 PubMed.

(10) Kalluri, R.; LeBleu, V. S. The biology, function, and biomedical applications of exosomes. Science 2020, 367 (6478). DOI: 10.1126/science.aau6977 PubMed. Lai, J. J.; Chau, Z. L.; Chen, S.-Y.; Hill, J. J.; Korpany, K. V.; Liang, N.-W.; Lin, L.-H.; Lin, Y.-H.; Liu, J. K.; Liu, Y.-C.; et al. Exosome Processing and Characterization Approaches for Research and Technology Development. Adv Sci (Weinh) 2022, 9 (15), e2103222. DOI: 10.1002/advs.202103222 PubMed. Wortzel, I.; Dror, S.; Kenific, C. M.; Lyden, D. Exosome-Mediated Metastasis: Communication from a Distance. Dev Cell 2019, 49 (3), 347–360. DOI: 10.1016/j.devcel.2019.04.011 PubMed. Yang, D.; Zhang, W.; Zhang, H.; Zhang, F.; Chen, L.; Ma, L.; Larcher, L. M.; Chen, S.; Liu, N.; Zhao, Q.; et al. Progress, opportunity, and perspective on exosome isolation - efforts for efficient exosome-based theranostics. Theranostics 2020, 10 (8), 3684–3707. DOI: 10.7150/thno.41580 PubMed. Wang, K.; Ye, H.; Zhang, X.; Wang, X.; Yang, B.; Luo, C.; Zhao, Z.; Zhao, J.; Lu, Q.; Zhang, H.; et al. An exosome-like programmable-bioactivating paclitaxel prodrug nanoplatform for enhanced breast cancer metastasis inhibition. Biomaterials 2020, 257. DOI: 10.1016/j.biomaterials.2020.120224. O’Brien, K.; Breyne, K.; Ughetto, S.; Laurent, L. C.; Breakefield, X. O. RNA delivery by extracellular vesicles in mammalian cells and its applications. Nat Rev Mol Cell Biol 2020, 21 (10), 585–606. DOI: 10.1038/s41580–020-0251-y PubMed. Li, S.-P.; Lin, Z.-X.; Jiang, X.-Y.; Yu, X.-Y. Exosomal cargo-loading and synthetic exosome-mimics as potential therapeutic tools. Acta Pharmacologica Sinica 2018, 39 (4), 542–551. DOI: 10.1038/aps.2017.178 PubMed.

(11) Llovet, J. M.; Willoughby, C. E.; Singal, A. G.; Greten, T. F.; Heikenwalder, M.; El-Serag, H. B.; Finn, R. S.; Friedman, S. L. Nonalcoholic steatohepatitis-related hepatocellular carcinoma: pathogenesis and treatment. Nat Rev Gastroenterol Hepatol 2023, 20 (8), 487–503. DOI: 10.1038/s41575–023-00754–7 From NLM Medline. Polyzos, S. A.; Kountouras, J.; Mantzoros, C. S. Obesity and nonalcoholic fatty liver disease: From pathophysiology to therapeutics. Metabolism 2019, 92, 82–97. DOI: 10.1016/j.metabol.2018.11.014 From NLM Medline.

(12) Rajesh, Y.; Sarkar, D. Molecular Mechanisms Regulating Obesity-Associated Hepatocellular Carcinoma. Cancers (Basel) 2020, 12 (5). DOI: 10.3390/cancers12051290 From NLM PubMed-not-MEDLINE. Clement, E.; Lazar, I.; Attané, C.; Carrié, L.; Dauvillier, S.; DucouxLJPetit, M.; Esteve, D.; Menneteau, T.; Moutahir, M.; Le Gonidec, S.; et al. Adipocyte extracellular vesicles carry enzymes and fatty acids that stimulate mitochondrial metabolism and remodeling in tumor cells. The EMBO Journal 2020, 39 (3). DOI: 10.15252/embj.2019102525. Li, X.; Li, C.; Zhang, L.; Wu, M.; Cao, K.; Jiang, F.; Chen, D.; Li, N.; Li, W. The significance of exosomes in the development and treatment of hepatocellular carcinoma. Mol Cancer 2020, 19 (1), 1. DOI: 10.1186/s12943–019-1085–0 PubMed.

(13) Lin, Y.-S.; Chen, W.-Y.; Liang, W.-Z. Investigation of Cytotoxicity and Oxidative Stress Induced by the Pyrethroid Bioallethrin in Human Glioblastoma Cells: The Protective Effect of Vitamin E (VE) and Its Underlying Mechanism. Chemical Research in Toxicology 2022, 35 (5), 880–889. DOI: 10.1021/acs.chemrestox.2c00033.

(14) Tian, C.; Guo, J.; Miao, Y.; Zheng, S.; Sun, B.; Sun, M.; Ye, Q.; Liu, W.; Zhou, S.; Kamei, K. I.;, et al. Triglyceride-Mimetic Structure-Gated Prodrug Nanoparticles for Smart Cancer Therapy. J Med Chem 2021, 64 (21), 15936–15948. DOI: 10.1021/acs.jmedchem.1c01328 From NLM Medline.

(15) Savina, A.; Furlan, M.; Vidal, M.; Colombo, M. I. Exosome release is regulated by a calcium-dependent mechanism in K562 cells. J Biol Chem 2003, 278 (22), 20083–20090. DOI: 10.1074/jbc.M301642200 From NLM Medline. Zhang, H.; Deng, T.; Liu, R.; Ning, T.; Yang, H.; Liu, D.; Zhang, Q.; Lin, D.; Ge, S.; Bai, M.; et al. CAF secreted miR-522 suppresses ferroptosis and promotes acquired chemo-resistance in gastric cancer. Mol Cancer 2020, 19 (1), 43. DOI: 10.1186/s12943–020-01168–8 From NLM Medline. Hu, J. L.; Wang, W.; Lan, X. L.; Zeng, Z. C.; Liang, Y. S.; Yan, Y. R.; Song, F. Y.; Wang, F. F.; Zhu, X. H.; Liao, W. J.; et al. CAFs secreted exosomes promote metastasis and chemotherapy resistance by enhancing cell stemness and epithelial-mesenchymal transition in colorectal cancer. Mol Cancer 2019, 18 (1), 91. DOI: 10.1186/s12943–019-1019-x From NLM Medline.

(16) Liu, X.; Cao, Z.; Wang, W.; Zou, C.; Wang, Y.; Pan, L.; Jia, B.; Zhang, K.; Zhang, W.; Li, W.;, et al. Engineered Extracellular Vesicle-Delivered CRISPR/Cas9 for Radiotherapy Sensitization of Glioblastoma. ACS Nano 2023, 17 (17), 16432–16447. DOI: 10.1021/acsnano.2c12857 PubMed. Jeppesen, D. K.; Fenix, A. M.; Franklin, J. L.; Higginbotham, J. N.; Zhang, Q.; Zimmerman, L. J.; Liebler, D. C.; Ping, J.; Liu, Q.; Evans, R.; et al. Reassessment of Exosome Composition. Cell 2019, 177 (2). DOI: 10.1016/j.cell.2019.02.029 PubMed.

(17) Hao, J.-W.; Wang, J.; Guo, H.; Zhao, Y.-Y.; Sun, H.-H.; Li, Y.-F.; Lai, X.-Y.; Zhao, N.; Wang, X.; Xie, C.;, et al. CD36 facilitates fatty acid uptake by dynamic palmitoylation-regulated endocytosis. Nature Communications 2020, 11 (1). DOI: 10.1038/s41467–020-18565–8.

(18) Luo, C.; Sun, J.; Sun, B.; He, Z. Prodrug-based nanoparticulate drug delivery strategies for cancer therapy. Trends Pharmacol Sci 2014, 35 (11), 556–566. DOI: 10.1016/j.tips.2014.09.008 PubMed. Zaidi, N.; Lupien, L.; Kuemmerle, N. B.; Kinlaw, W. B.; Swinnen, J. V.; Smans, K. Lipogenesis and lipolysis: the pathways exploited by the cancer cells to acquire fatty acids. Prog Lipid Res 2013, 52 (4), 585–589. DOI: 10.1016/j.plipres.2013.08.005 PubMed. Zechner, R.; Zimmermann, R.; Eichmann, T. O.; Kohlwein, S. D.; Haemmerle, G.; Lass, A.; Madeo, F. FAT SIGNALS--lipases and lipolysis in lipid metabolism and signaling. Cell Metabolism 2012, 15 (3), 279–291. DOI: 10.1016/j.cmet.2011.12.018 PubMed.

(19) Zhu, H.; Zhou, W.; Wan, Y.; Lu, J.; Ge, K.; Jia, C. Light-activatable multifunctional paclitaxel nanoprodrug for synergistic chemo-photodynamic therapy in liver cancer. Biofactors 2022, 48 (4), 918–925. DOI: 10.1002/biof.1832 PubMed.

(20) Mulcahy, L. A.; Pink, R. C.; Carter, D. R. F. Routes and mechanisms of extracellular vesicle uptake. Journal of Extracellular Vesicles 2014, 3 (1). DOI: 10.3402/jev.v3.24641.

(21) Peche, V. S.; Pietka, T. A.; Jacome-Sosa, M.; Samovski, D.; Palacios, H.; Chatterjee-Basu, G.; Dudley, A. C.; Beatty, W.; Meyer, G. A.; Goldberg, I. J.;, et al. Endothelial cell CD36 regulates membrane ceramide formation, exosome fatty acid transfer and circulating fatty acid levels. Nature Communications 2023, 14 (1), 4029. DOI: 10.1038/s41467–023-39752–3 PubMed.

(22) Yan, C.; Tian, X.; Li, J.; Liu, D.; Ye, D.; Xie, Z.; Han, Y.; Zou, M.-H. A High-Fat Diet Attenuates AMPK α1 in Adipocytes to Induce Exosome Shedding and Nonalcoholic Fatty Liver Development In Vivo. Diabetes 2021, 70 (2), 577–588. DOI: 10.2337/db20–0146 PubMed.

(23) Sivasami, P.; Elkins, C.; Diaz-Saldana, P. P.; Goss, K.; Peng, A.; Hamersky, M.; Bae, J.; Xu, M.; Pollack, B. P.; Horwitz, E. M.;, et al. Obesity-induced dysregulation of skin-resident PPARγ+ Treg cells promotes IL-17A-mediated psoriatic inflammation. Immunity 2023, 56 (8). DOI: 10.1016/j.immuni.2023.06.021 PubMed.

(24) Tian, C.; Guo, J.; Miao, Y.; Zheng, S.; Sun, B.; Sun, M.; Ye, Q.; Liu, W.; Zhou, S.; Kamei, K.-i.;, et al. Triglyceride-Mimetic Structure-Gated Prodrug Nanoparticles for Smart Cancer Therapy. Journal of Medicinal Chemistry 2021, 64 (21), 15936–15948. DOI: 10.1021/acs.jmedchem.1c01328.

(25) Raza, A.; Rossi, G. R.; Janjua, T. I.; Souza-Fonseca-Guimaraes, F.; Popat, A. Nanobiomaterials to modulate natural killer cell responses for effective cancer immunotherapy. Trends Biotechnol 2023, 41 (1), 77–92. DOI: 10.1016/j.tibtech.2022.06.011 From NLM Medline. Huang, R.; Zhou, P.; Chen, B.; Zhu, Y.; Chen, X.; Min, Y. Stimuli-Responsive Nanoadjuvant Rejuvenates Robust Immune Responses to Sensitize Cancer Immunotherapy. ACS Nano 2023. DOI: 10.1021/acsnano.3c06233 PubMed. Samain, R.; Maiques, O.; Monger, J.; Lam, H.; Candido, J.; George, S.; Ferrari, N.; KohIhammer, L.; Lunetto, S.; Varela, A.; et al. CD73 controls Myosin II-driven invasion, metastasis, and immunosuppression in amoeboid pancreatic cancer cells. Sci Adv 2023, 9 (42), eadi0244. DOI: 10.1126/sciadv.adi0244 PubMed.

(26) Qi, R.; Sammler, E.; Gonzalez-Hunt, C. P.; Barraza, I.; Pena, N.; Rouanet, J. P.; Naaldijk, Y.; Goodson, S.; Fuzzati, M.; Blandini, F.;, et al. A blood-based marker of mitochondrial DNA damage in Parkinson’s disease. Sci Transl Med 2023, 15 (711), eabo1557. DOI: 10.1126/scitranslmed.abo1557 PubMed. Turner, L.; Martinez, J. R.; Najjar, S.; Rajapaksha Arachchilage, T.; Wang, J. C. Businesses marketing purported stem cell treatments and exosome therapies for COVID-19: An analysis of direct-to-consumer online advertising claims. Stem Cell Reports 2023. DOI: 10.1016/j.stemcr.2023.09.015 PubMed.

(27) Xu, Y.; Yao, Y.; Yu, L.; Fung, H. L.; Tang, A. H. N.; Ng, I. O.-L.; Wong, M. Y. M.; Che, C.-M.; Yun, J. P.; Cui, Y.;, et al. Clathrin light chain A facilitates small extracellular vesicle uptake to promote hepatocellular carcinoma progression. Hepatol Int 2023. DOI: 10.1007/s12072–023-10562–5 PubMed.

(28) Holmström, K. M.; Finkel, T. Cellular mechanisms and physiological consequences of redox-dependent signalling. Nat Rev Mol Cell Biol 2014, 15 (6), 411–421. DOI: 10.1038/nrm3801 PubMed.

(29) Wang, X.; Xia, J.; Yang, L.; Dai, J.; He, L. Recent progress in exosome research: isolation, characterization and clinical applications. Cancer Gene Ther 2023, 30 (8), 1051–1065. DOI: 10.1038/s41417–023-00617-y PubMed.

(30) Gunassekaran, G. R.; Poongkavithai Vadevoo, S. M.; Baek, M.-C.; Lee, B. M1 macrophage exosomes engineered to foster M1 polarization and target the IL-4 receptor inhibit tumor growth by reprogramming tumor-associated macrophages into M1-like macrophages. Biomaterials 2021, 278, 121137. DOI: 10.1016/j.biomaterials.2021.121137 PubMed.

