## Supplementary figures and images for "Hybrid Adipocyte-Derived Exosome Nano Platform for Potent Chemo-Phototherapy in Targeted Hepatocellular Carcinoma"

### (A) Mass spectrum and (B) 1H NMR spectrum in CDCl3 of DSTG.

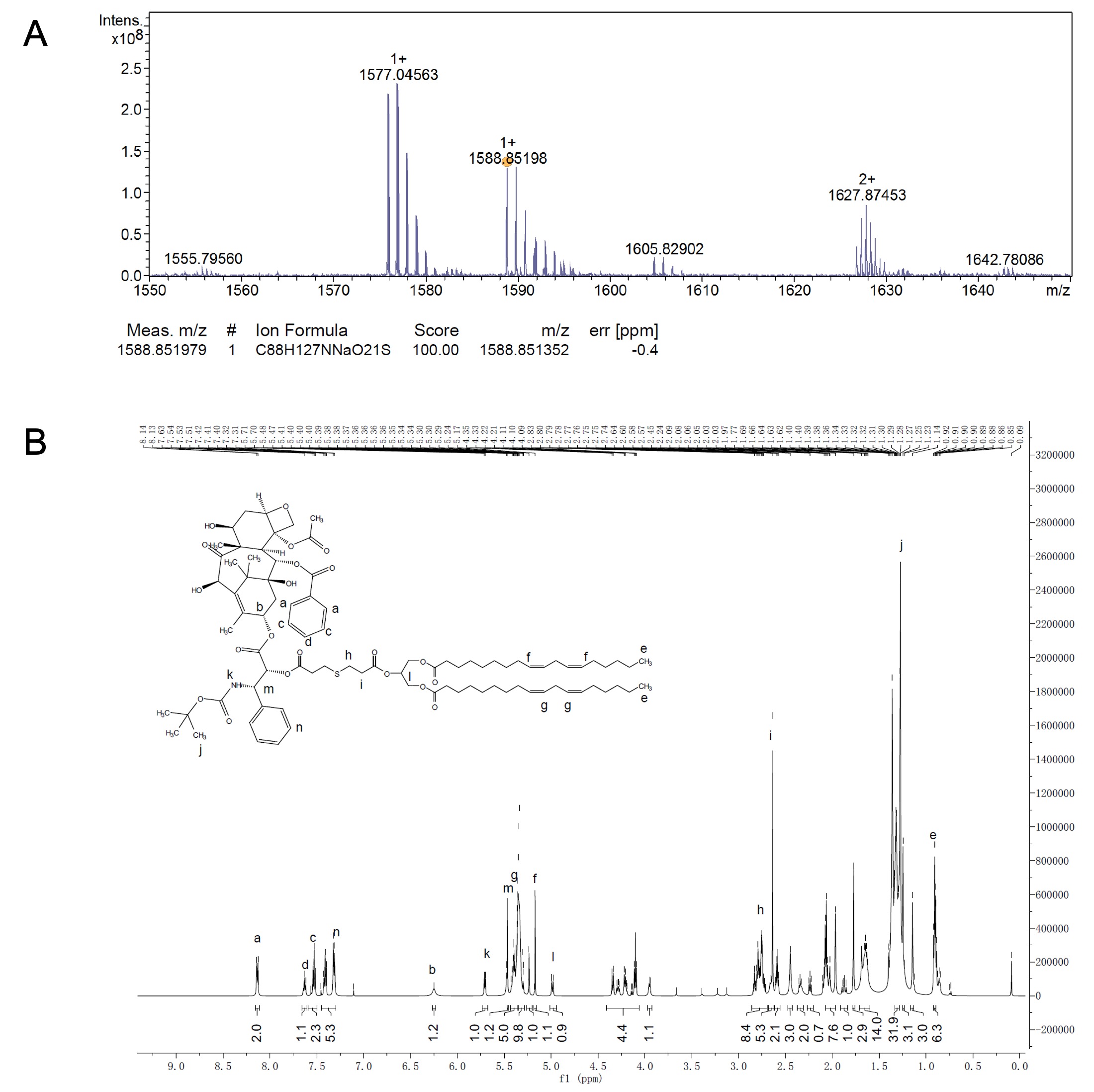

### (A) Mass spectrum and (B) 1H NMR spectrum in CDCl3 of PPLA.

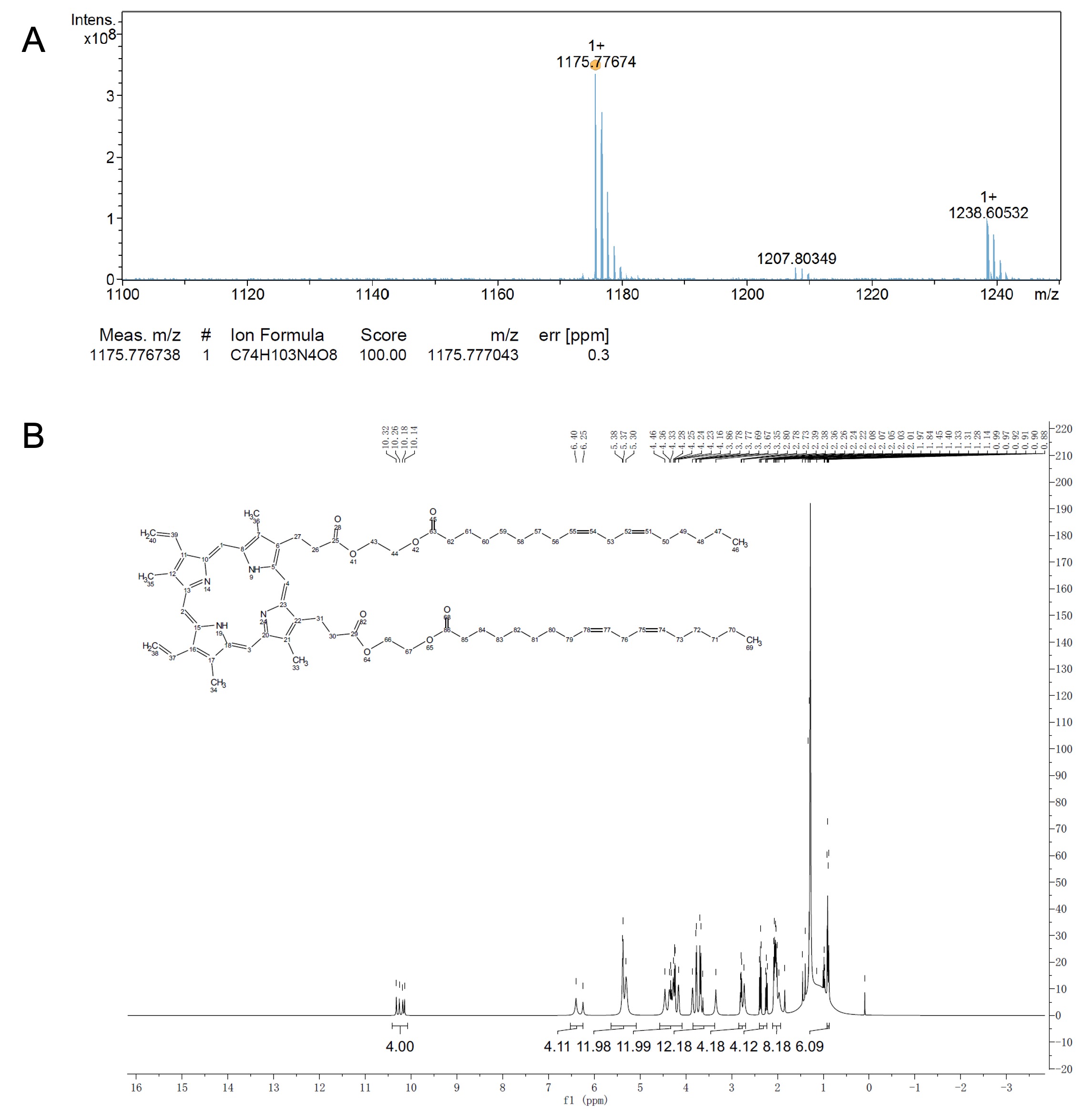

### (A)The fluorescence spectra and (B) ultraviolet spectra of PpIX and PPLA.

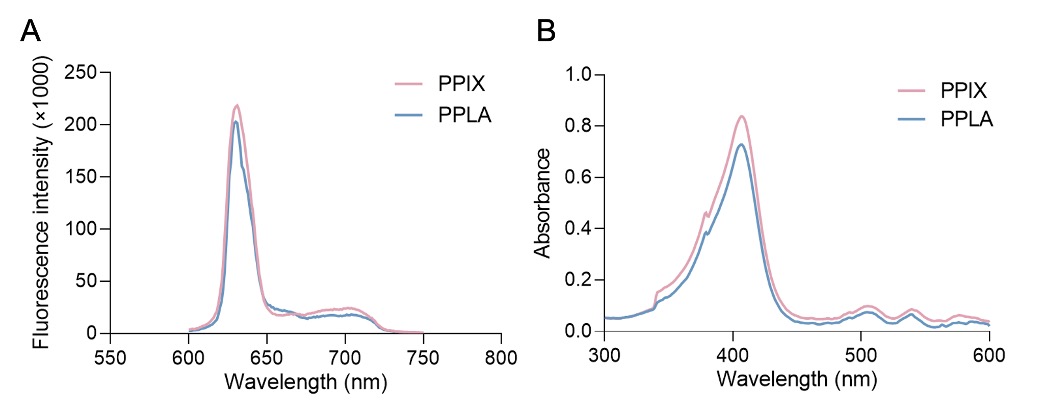

### Cellular uptake of HepG2 cancer cells.

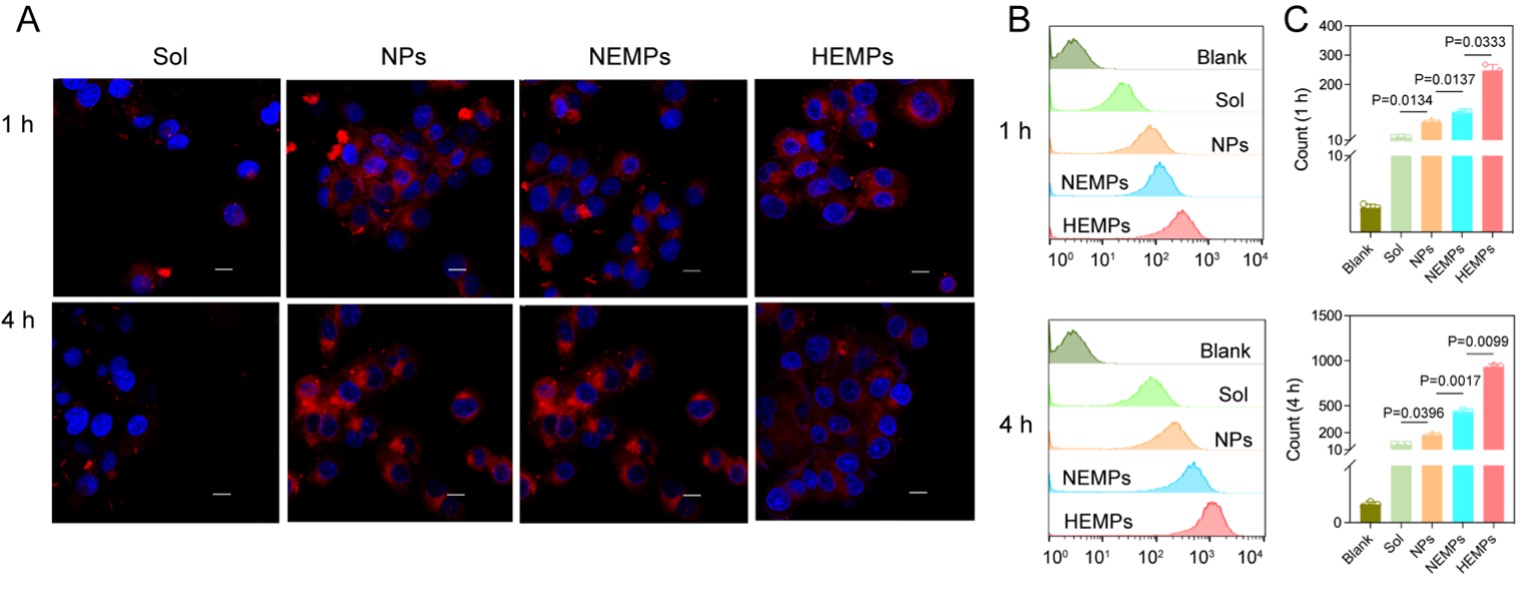

### Fluorescence distribution from flow cytometry of Hepa1-6 cells incubated for (A) 1 h or (B) 4 h.

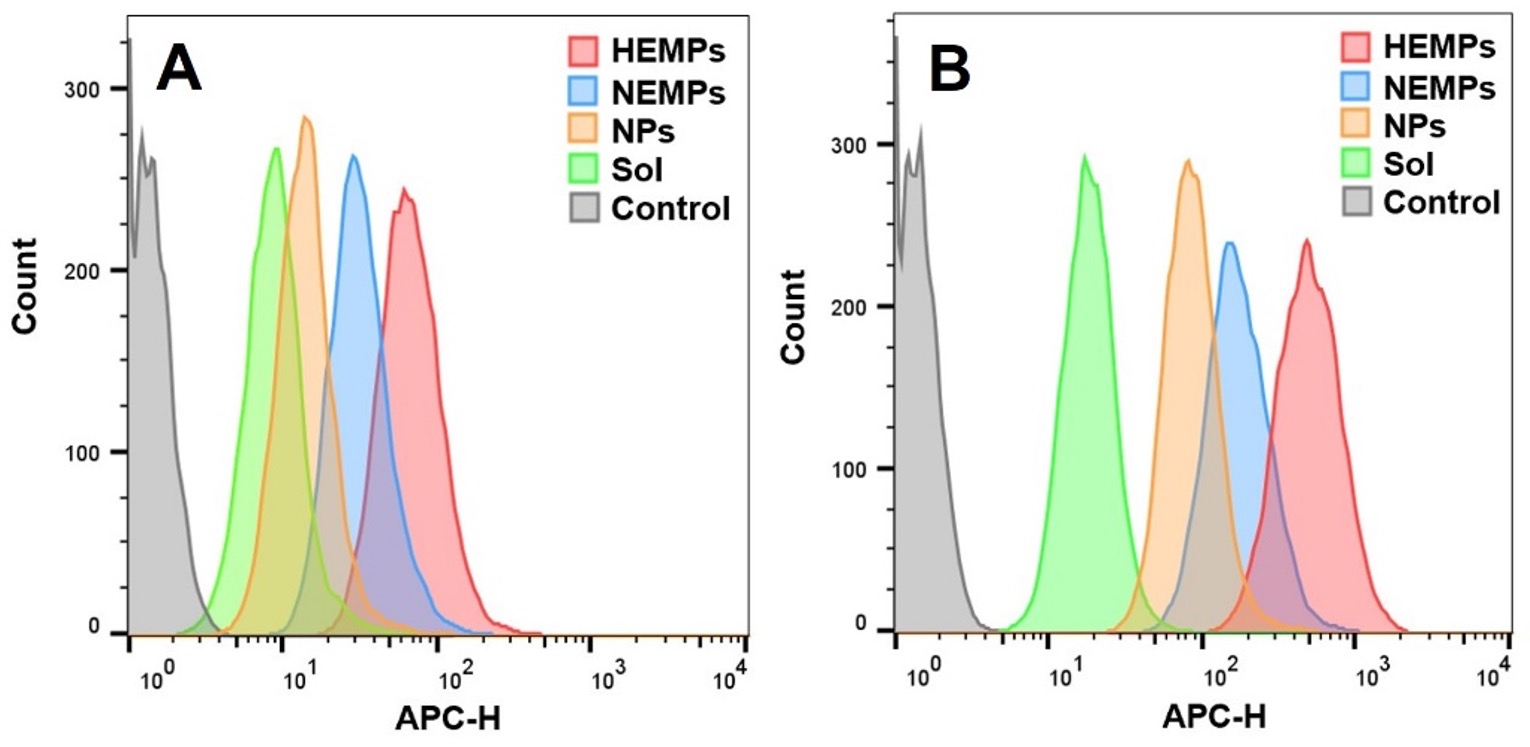

### In vitro cellular lipid peroxidation level.

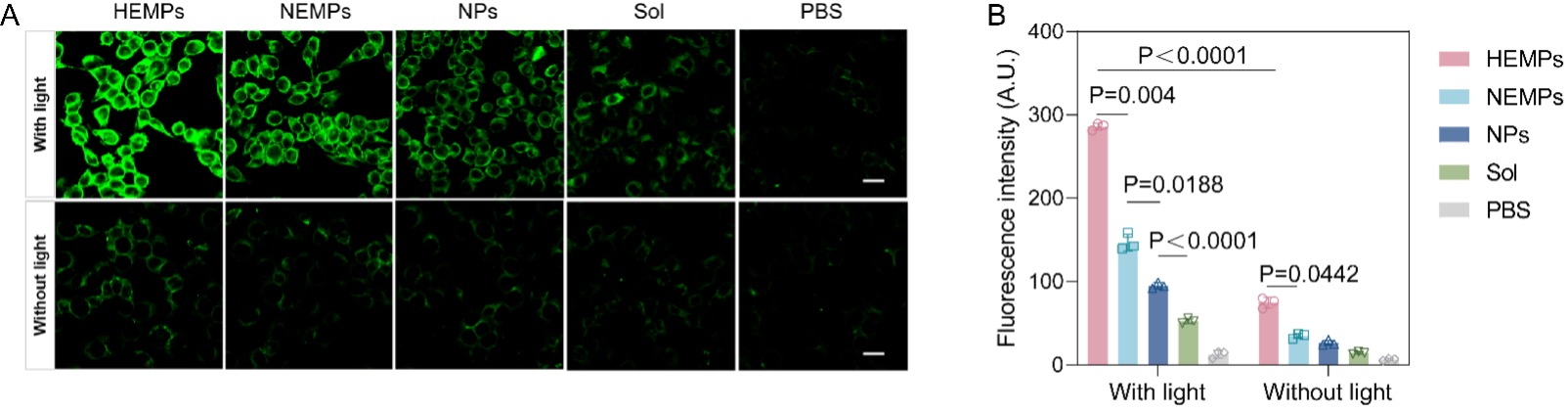

### In vitro cytotoxicity (IC50 values).

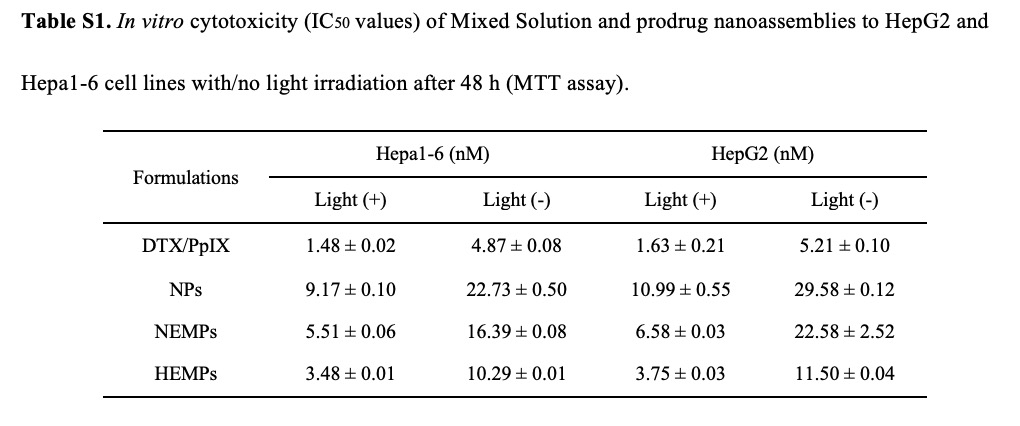

### Pharmacokinetic parameters of Mixed solution, HEMPs, NEMPs, and NPs.

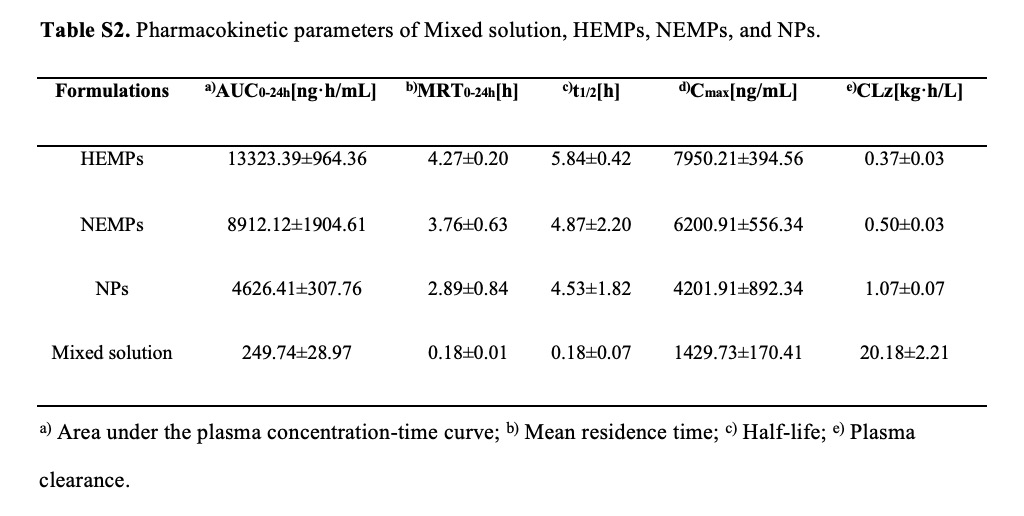

### Supplemental Data 1

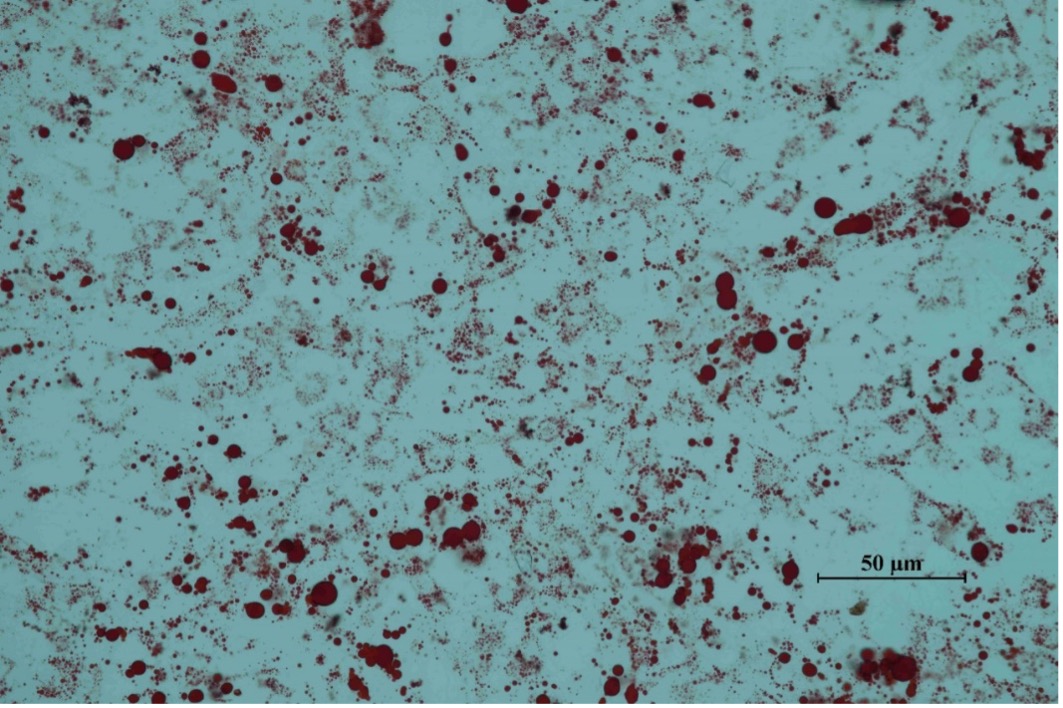

### Supplemental Data 2

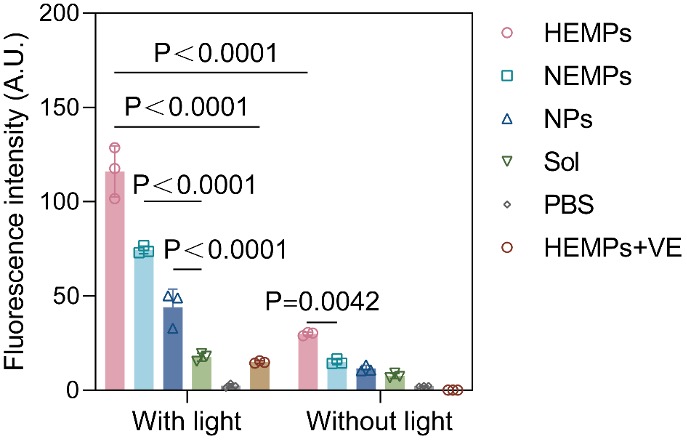

### Supplemental Data 3

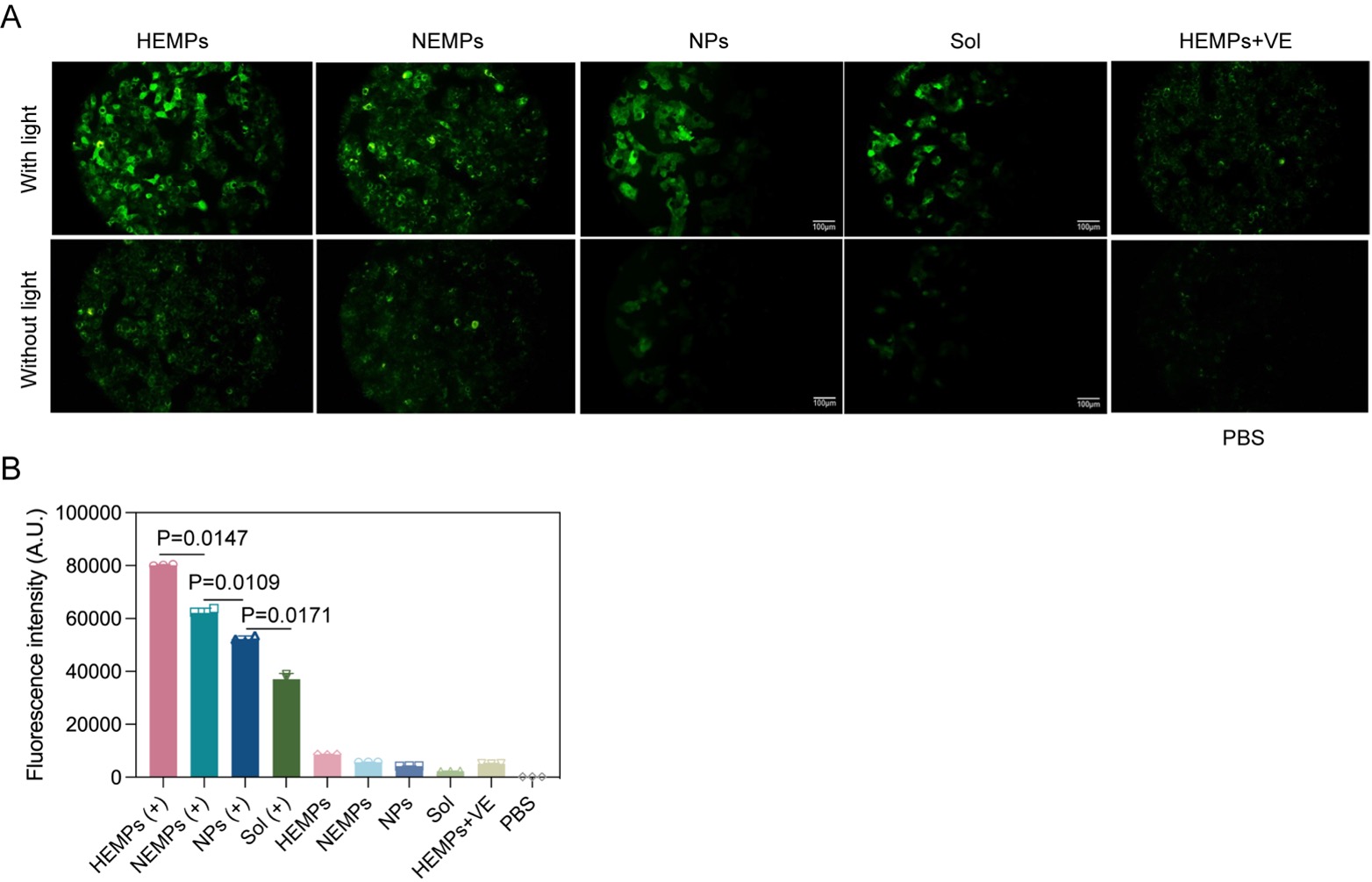

### Synthesis pathways of DSTG.

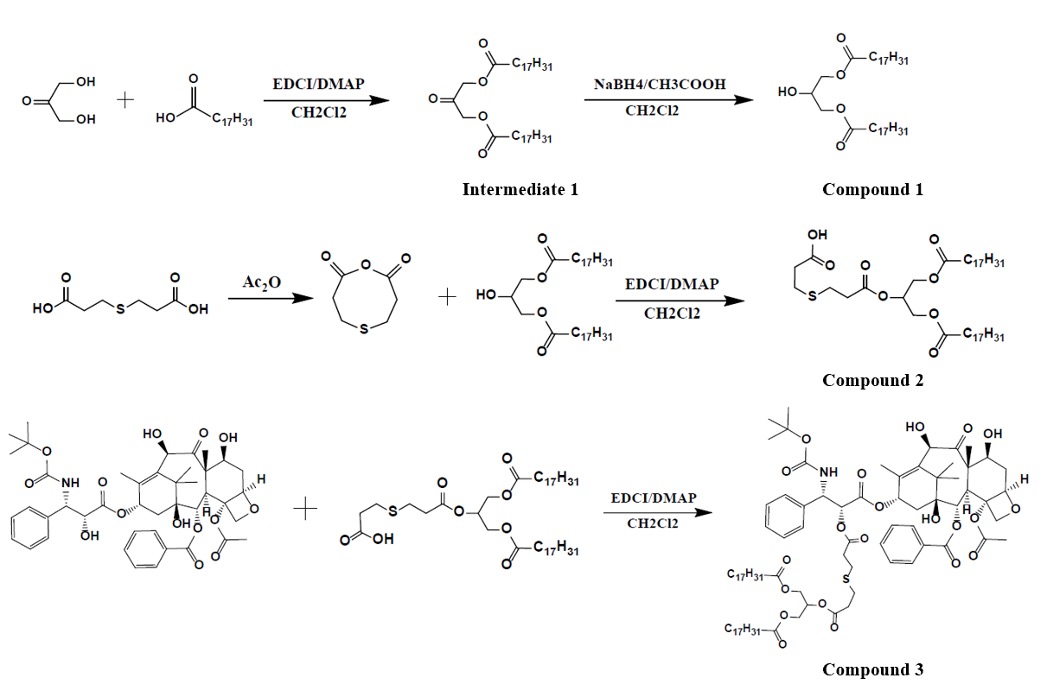

### Synthesis pathways of PPLA.

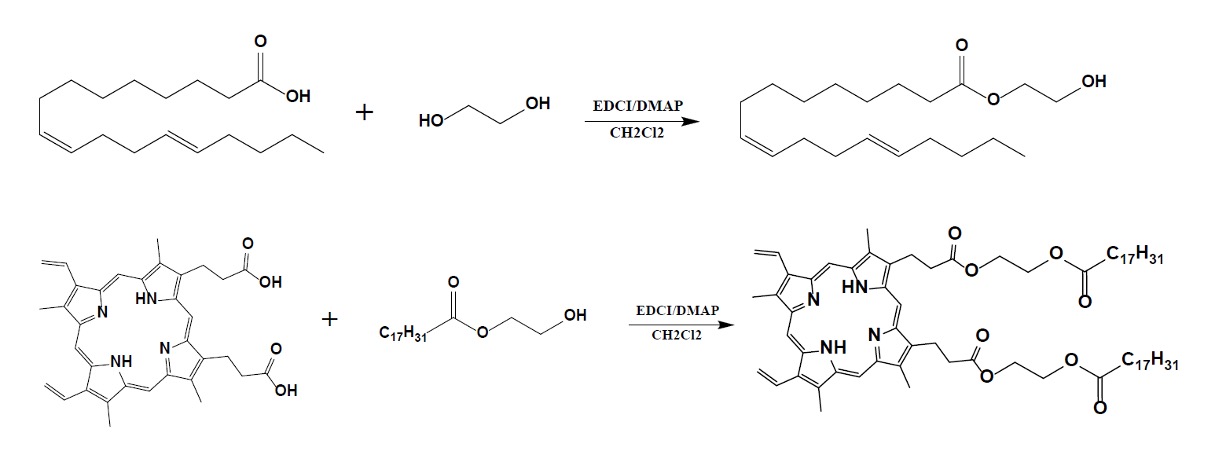

### The oxidative release mechanism of the prodrug nanoparticles.

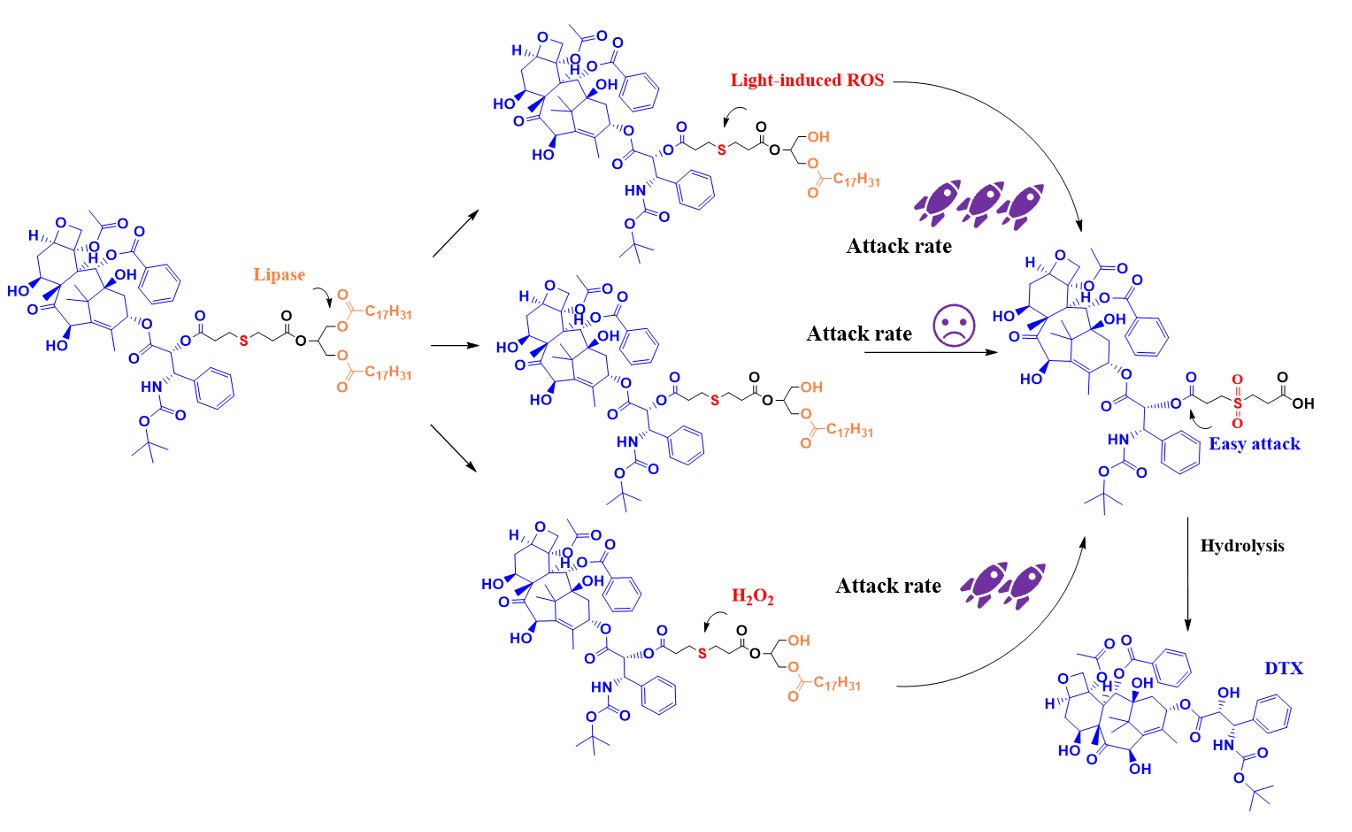

### Western blotting of characteristic markers CD36, CD9, CD81, TSG101, and GAPDH. (a: normal treatment; b: palmitic acid treatment)

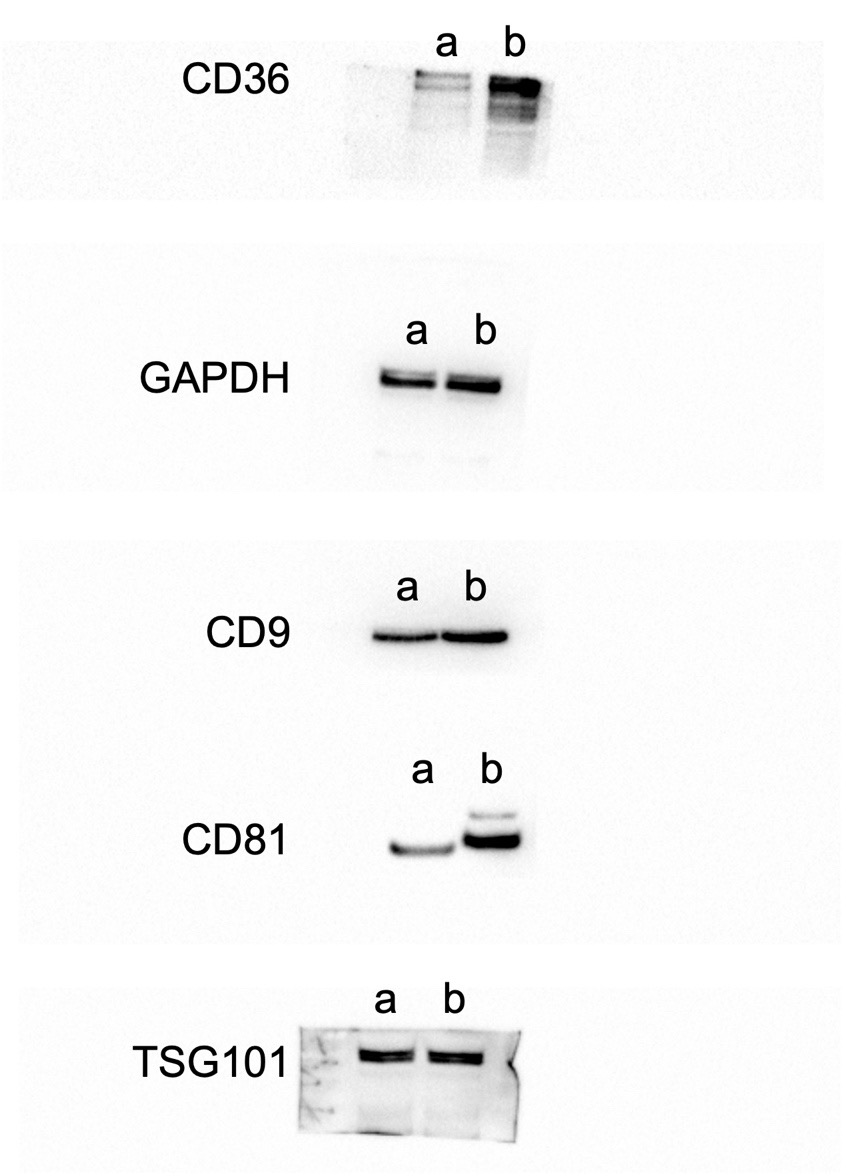
